# Role of auditory feedback for vocal production learning in the Egyptian fruit-bat

**DOI:** 10.1101/2023.11.21.568126

**Authors:** Julie E. Elie, Sandra E. Muroy, Daria Genzel, Tong Na, Lisa A. Beyer, Donald L. Swiderski, Yehoash Raphael, Michael M. Yartsev

## Abstract

Some species have evolved the ability to use the sense of hearing to modify existing vocalizations, or even create new ones. This ability corresponds to various forms of vocal production learning that are all possessed by humans, and independently displayed by distantly related vertebrates. Among mammals, a few species, including the Egyptian fruit-bat, would possess such vocal production learning abilities. Yet the necessity of an intact auditory system for the development of the Egyptian fruit-bat typical vocal repertoire has not been tested. Furthermore, a systematic causal examination of learned and innate aspects of the entire repertoire has never been performed in any vocal learner. Here we addressed these gaps by eliminating pups’ sense of hearing at birth and assessing its effects on vocal production in adulthood. The deafening treatment enabled us to both causally test these bats vocal learning ability and discern learned from innate aspects of their vocalizations. Leveraging wireless individual audio recordings from freely interacting adults, we show that a subset of the Egyptian fruit-bat vocal repertoire necessitates auditory feedback. Intriguingly, these affected vocalizations belong to different acoustic groups in the vocal repertoire of males and females. These findings open the possibilities for targeted studies of the mammalian neural circuits that enable sexually dimorphic forms of vocal learning.

## Introduction

Vocalizations constitute the building blocks of the acoustic code used for vocal communication. The efficiency and complexity of a vocal communication system depends on the species ability to produce a diversity of such building blocks. While this diversity can be supported by hard wired motor programs that are innate (e.g., motor programs of male mouse song^1^), the sound diversity is also enhanced in some species by their ability to modify existing calls or create new ones^2–7^. This ability is defined as vocal production learning and has been examined in a wide variety of species^2,4,8^. Yet, experiments discerning innate versus learned vocalizations in the entire repertoire of a single species are missing.

The Egyptian fruit-bat has been proposed as one of the species capable of vocal production learning after several studies demonstrated how manipulation of either the social or physical environment affects the acoustic characteristics of their communication calls^9–11^. However, it remains unknown whether all or only a subset of their repertoire is acquired through learning. Furthermore, the acoustic characteristics of the voice of individuals can easily be mistaken for an effect of vocal learning when comparing the vocal behaviors of two groups of animals (one control and one experimental). Previous conclusions on vocal learning abilities are weakened by the lack of reliable identification of vocalizing individuals^9–11^. Combined, these gaps limit thorough exploration of the neural mechanisms of vocal production learning in this species, although cutting-edge technologies for measuring brain activity are readily available^12–15^

Here we reasoned that because vocal production learning heavily depends on the sense of hearing, the main sensory feedback that the nervous system can use to change or create new motor programs^16^, abolishing this feedback during development can begin revealing what aspects of the repertoire are learned and which are innate. Therefore, we conducted the first causal study of vocal production learning in the Egyptian fruit-bat by establishing a method for deafening pups shortly after birth and testing the impact of such manipulation on the vocal behavior of each of these bats when they became adults. Because the ontogeny and maturation of the Egyptian fruit-bat vocal behavior is unknown, we focused our investigation of the vocalizations of hearing and deaf bats when they were fully mature adults (Figure 1A and 1D, Methods). By then, the bats had lived for years in a communal social environment where they were equally exposed to conspecific vocalizations. Over a seven week period, the deaf and hearing adults were co-housed and daily equipped with wireless audiologger collars, that enabled to detect all vocalizations from the bat repertoire and to unambiguously assign the vocalizer identity to each vocalization^13^. Additionally, an ambient microphone further ensured high quality audio recordings of the vocalizations. After manual curation, we obtained thousands of calls from both hearing (n=16,937) and deaf bats (n=11,154).

## Results

### Deafening at birth and recording of vocalizations at adulthood

The acoustic isolation of newborn pups to any bat vocalization was ensured by single-housing pregnant mothers before pup birth in acoustically isolated chambers. Deafening of the pups was then chemically achieved shortly after birth (Figure 1A; n=10 bats). Five pups received an injection regimen of the deafening antibiotic kanamycin, while five pups received control saline injections (Methods). In each experimental group of five, there were three females and two males. The extent of hearing loss incurred was verified both functionally, using auditory brainstem recordings (Figure 1B, Figure S1; Methods), and histologically, by examining the cochlear epithelium (Figure 1C, Figure S1; Methods). These approaches revealed profound deafness in the kanamycin treated bats and normal hearing in the saline treated bats.

**Figure 1.**
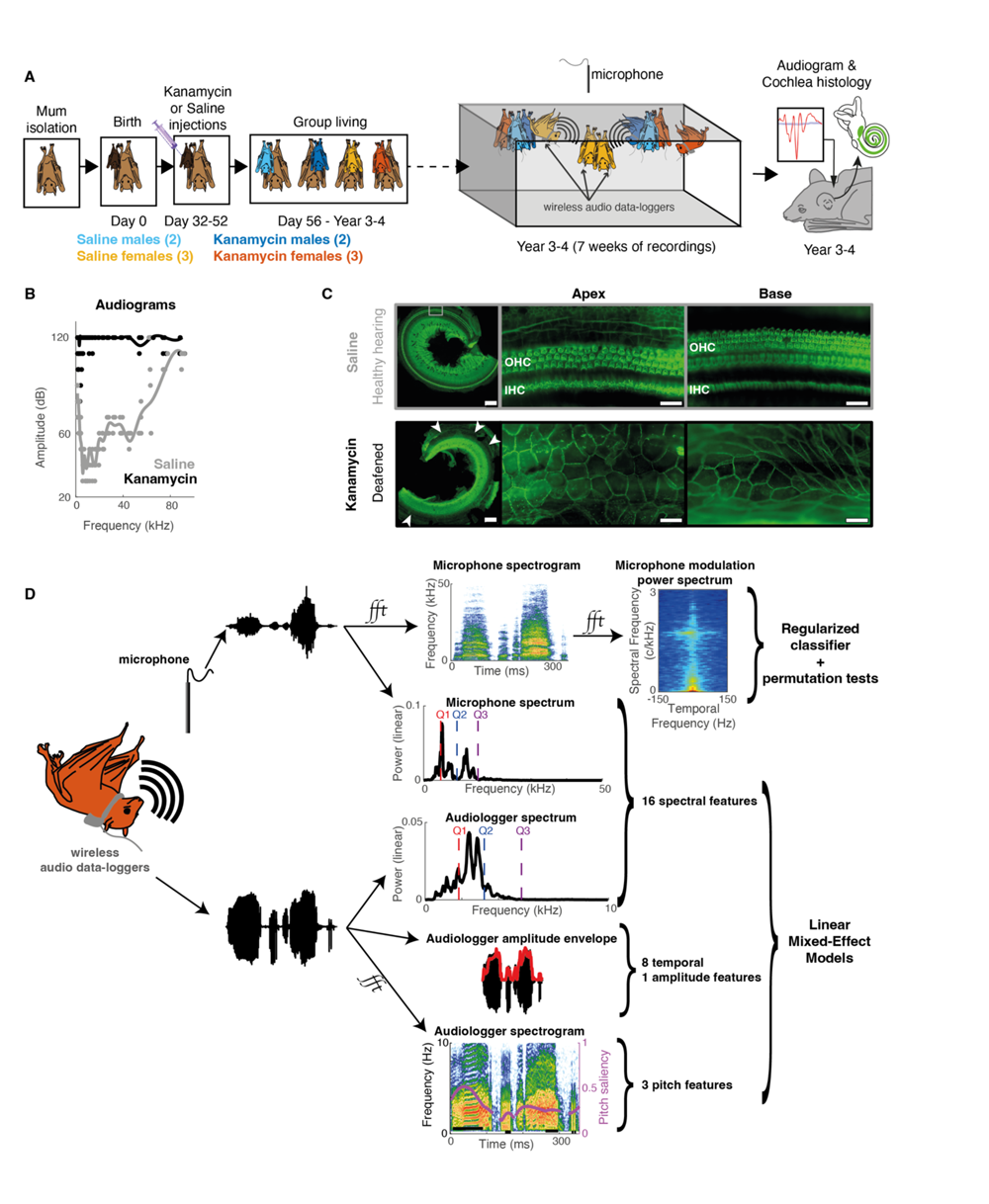
Experimental design testing the necessity of hearing from birth for the normal development of the vocal repertoire in adult Egyptian fruit-bats. A. Pregnant females gave birth and cared for their young in complete isolation from the rest of the colony for two months. Their ten pups were randomly assigned to a treatment by kanamycin or saline (control). Mothers and pups were then reunited with the rest of the colony for 3-4 years. Well after reaching adulthood, the ten manipulated bats were housed together and their adult vocal behavior was recorded for seven weeks. Finally, their audiograms were measured and their cochlea inspected for histological damage. B. Audiograms obtained from measuring the Auditory Brainstem Response of all bats (n=10) to tone pips covering the species auditory range. Dots depict individual values, while lines show the average fitted with a spline. Note that responses of the auditory nerve were absent (values ceiled at 120dB) or only detectable for very loud sounds (>90dB) in kanamycin treated bats, while the expected audiogram was obtained for saline treated bats. C. Representative examples of the hearing sensory epithelium (organ of Corti) at two levels of the cochlea (apical turn, first and second columns; basal turn, third column) in two bats stained with phalloidin (first row, female; second row, male). Scale bars: 200µm in first column, 25µm in other columns; grey square: area shown in second column. Note the intact epithelium (OHC, outer hair cells; IHC, inner hair cells) both in the apical and basal cochlear areas of the saline bat, as opposed to the non-sensory flat epithelium in the kanamycin bat. Hair cells are absent throughout the organ of Corti (see white arrows) except for a few cells in kanamycin female bats (Figure S1). D. Vocalizations are recorded both from an environment microphone and a piezo sensor placed against the throat of the bat (wireless audiologger). Two approaches are used to find the effect of deafening on bats’ vocalizations. Upper panel: the data driven approach is using the modulation power spectrum of vocalizations as input to a classifier that is trained and tested in cross-validation to discriminate between hearing and deaf bats’ calls. The permutation test gives the correct chance level taking into account any acoustic differences due to bat identity. The second approach test the effect of deafening by fitting a Linear Mixed-Effect model controlling for the identity of the bats on each of 28 acoustic features of the vocalizations (Figure S3 and methods).

### No effect of deafening on the whole repertoire when controlling for identity

Deafening did not affect the vocal activity of bats (Figure S2) and we obtained a similar number of vocalizations from both deafened and hearing bats (>1,000 vocalizations on average per individual). To explore if deafening was affecting the acoustic features of vocalizations, we used both a classical bioacoustical approach (28 predefined acoustic features measured on each call) and a data driven approach (Figure 1D). In the latter approach, we used the Modulation Power Spectrum^17^ (MPS) to describe the spectro-temporal modulations of vocalizations. The resulting MPS of each vocalization was used both in UMAP projections of all vocalizations and in regularized linear classifiers that quantified the level of discrimination between calls emitted by deaf and hearing bats (Regularized Permutation Discriminant Function Analysis, RPDFA, see Methods). Unexpectedly, with regards to previous reports^9–11^, when we considered all the recorded vocalizations, none of the two approaches revealed any effect of deafness when the identity of the vocalizing bats was accounted for (Figure S3, all 28 linear mixed-effects models on acoustic features with p>0.05). The performance of the RPDFA at classifying call MPS between deaf and hearing bats was 60.4% ± 1% and not significantly different from the chance level of classifying any random two groups of five bats, 62.4% ± 1.3% (Figure S4A and S4B; note that considering the voice characteristics of individuals raises the chance level higher than 50%).

### Heterogeneity in vocal space coverage according to sex and hearing ability

However, we next considered the possibility that not all vocalizations necessarily had to depend on auditory feedback. That is, some could be innate whereas others could be learned. Therefore, considering all vocalizations together, as was done in previous studies, may result in masking the effect of deafening. Indeed, linear measures of acoustic proximity revealed that bats were not homogeneously occupying the vocal space (Figure 2). First, we found that males covered a larger vocal space than females (vocalizations with large values of Sex Dissimilarity; Figure 2A and Figure S5A, S5B and S5D) and that for each sex, some vocalizations were distinct between deaf and hearing bats (Figure S6A-6C; performance of the RPDFA against identity controlled chance level: all male calls, 67.4 ± 1.2% vs 64.5 ± 0.8%; all female calls, 62.0 ± 0.6% vs 58.6 ± 1.3%). Second, a small proportion of the repertoire of each sex was idiosyncratic to the hearing ability of bats (Figure 2B and 2D). In males, these distinctive vocalizations represented 23.9 ± 3.2% of hearing bats repertoire and could not be matched with similar calls from deaf bats, indicating that hearing males produced vocalizations that deaf males did not (Figure 2B, Figure S7). In females, the few distinctive vocalizations represented 4.5 ± 1.5% of deaf bats repertoire and did not have a match with vocalizations produced by hearing females, indicating that these deaf females were producing a few atypical vocalizations (Figure 2D, Figure S7). These vocalizations belonged to regions of the vocal space that grouped the loudest, more tonal, and higher pitch vocalizations in the repertoire (Figure S8).

**Figure 2.**
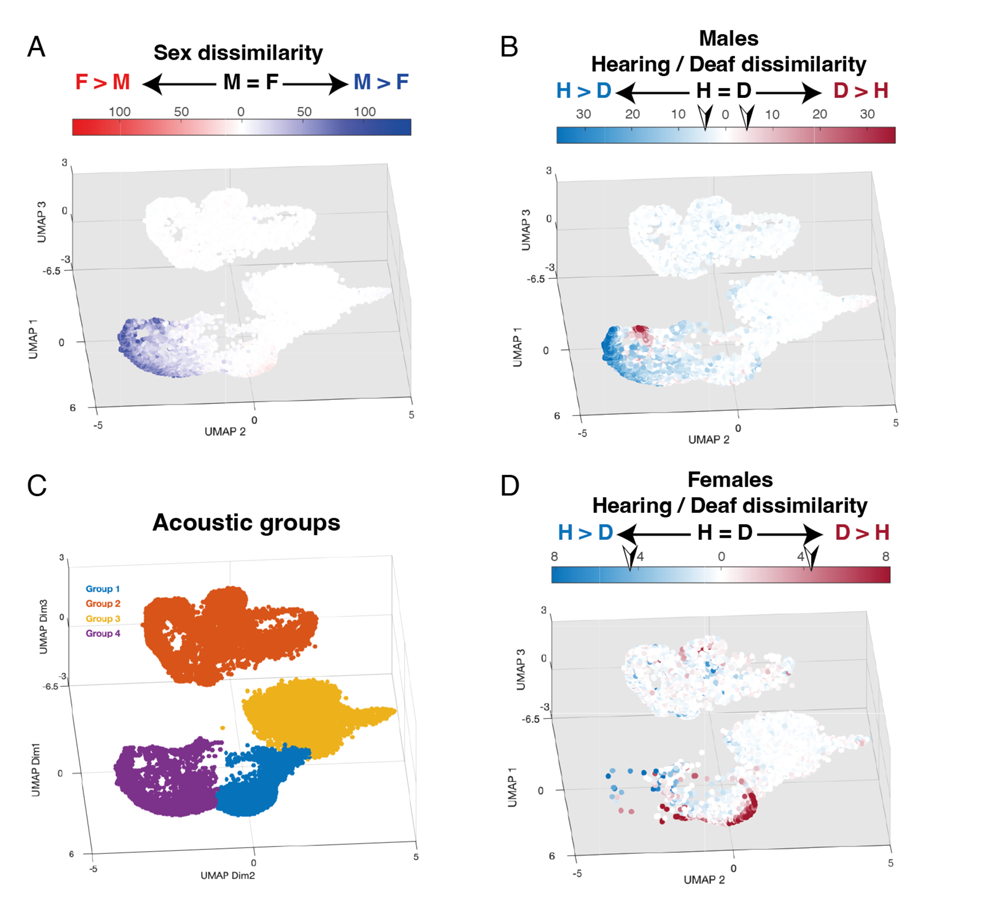
Effect of sex and deafness through the vocal space. Projections of vocalizations as represented by their MPS in a 3-dimension UMAP, the vocal space. A. Dissimilarity of bat vocalizations to opposite sex vocalizations. The color of the points indicate the sex of the emitter (male, blue, and female, red) and the intensity of the color the linear PCA distance to the closest ten vocalizations produced by an opposite sex bat (see methods). Note that vocalizations with UMAP2<-2 are almost exclusively produced by males. B and D Dissimilarity of male (B) or female (D) vocalizations to vocalizations from a same-sex bat with opposite hearing ability. Male (B) or female (D) vocalizations are color coded according to the hearing ability of the emitter (hearing, blue, and deaf, red). The intensity of the color corresponds to the linear PCA distance to the closest ten vocalizations produced by a male (B) or a female (D) of opposite hearing ability (methods). The arrowhead on the color scale indicates the threshold over which vocalizations were considered idiosyncratic of the bat hearing ability (dissimilarity value larger than expected on average to any same sex calls; see Figure S8 and methods). C. Parsing of the vocal space into four groups according to the results of a Hierarchical Agglomerative Clustering algorithm.

To further quantify and identify the spectro-temporal differences between deaf and hearing bats’ vocalizations throughout the repertoire, we parsed the vocal space into four groups (Figure 2C, Figure S9). This exact number of groups optimized both maximum division and cluster quality (Figure S10). Deafness affected vocalizations in two of the four groups (Figure S4; performance of the RPDFA against the identity-controlled chance level: Group 1, 73.2 ± 3.7% vs 66.8 ± 3.2%; Group 4, 85.9 ± 1.8% vs 76.9 ± 2.5%). Group 4 corresponded to loud and high pitch vocalizations that females rarely produced (Figure 2A, Figure S5C, S8 and S9) and only male calls were different between hearing and deaf individuals in this acoustic group (Figure S6G-6I; performance of the RPDFA against identity controlled chance level: Male Group 4 calls, 89.5 ± 2.7% vs 79.2 ± 2.1%; Female Group 4, 62 ±10.4% vs 59.7 ± 9.2%). Surprisingly, although both sexes produced calls that belonged to acoustic Group 1 (Figure S5C), only female calls were distinct between hearing and deaf bats (Figure S6D-6F; performance of the RPDFA against identity controlled chance level: Male Group 1 calls, 69.9 ± 5.4% vs 65.7 ± 3.9%; Female Group 1, 76.4 ± 1.9% vs 61.2 ± 3.1%). Combined, these findings suggest that a subset of the Egyptian fruit bat repertoire depended on intact auditory feedback in a sexually dimorphic manner.

### Effect of deafening on the spectro-temporal patterns of a subset of male and female calls

The acoustic differences in the vocalizations of deaf and hearing bats were a complex combination of both spectral and temporal parameters. The atypical vocalizations from deaf females in acoustic group1 were characterized by complex patterns of pitch modulations and decreased formants power (Figure S11). However, these differences were not captured by the classical bioacoustical approach (Figure S12). In contrast, the pattern of spectro-temporal modulations that characterized hearing male calls from acoustic group 4 was clear. Hearing males produced simple downsweep calls that were never matched by deaf males who produced calls with more upsweeps (Figure 3, Figure S13). Furthermore, deaf male calls had higher spectral content (i.e. more harmonics) than hearing male calls as measured by spectral features such as the third spectral quartile (Figure 3, Figure S13).

**Figure 3.**
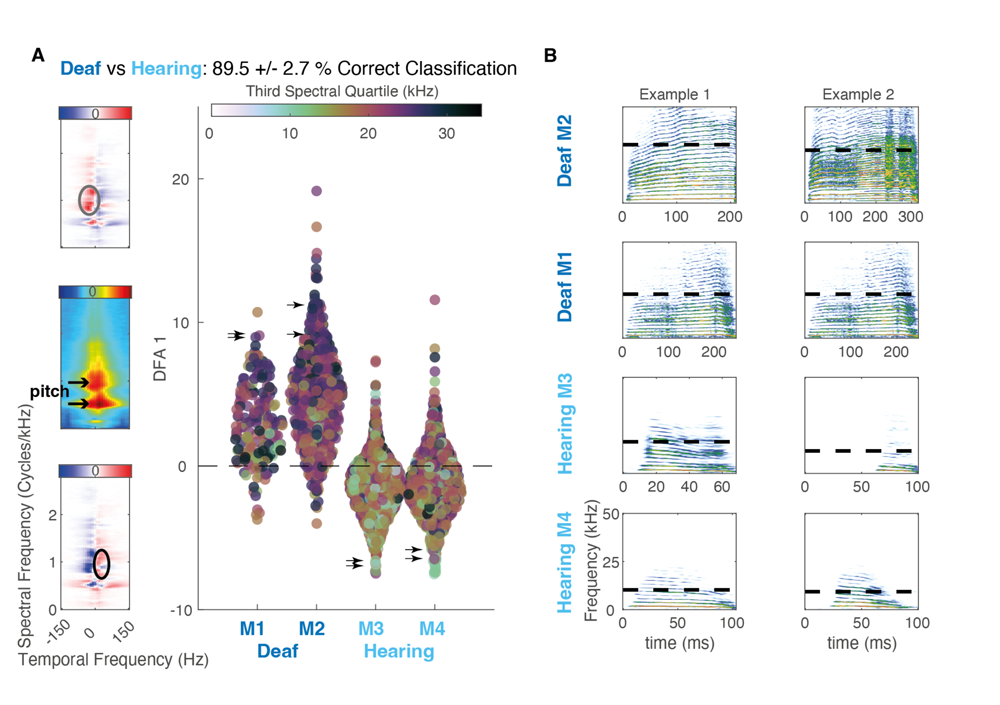
Effect of deafness on male calls in acoustic group 4. A. Coefficients of the acoustic Group 4 vocalizations of each male bat when projected on the first linear acoustic axis of the PCARDFA that best discriminates between deaf (M1 and M2) and hearing (M3 and M4) bats. Each circle in the scatter plot represents a vocalization. The circle color indicates the value of the Third Spectral Quartile of the vocalization. The three images on the left show the average MPS of male vocalizations in acoustic Group 4 (middle) and the spectro-temporal changes in the MPS (discriminant function) that best characterize vocalizations from deaf males (top) and vocalizations from hearing males (bottom). Note that while on average male vocalizations in acoustic Group 4 are characterized by pitch values of 1c/kHz or 0.5 c/kHz corresponding respectively to 1kHz and 2kHz (black arrows), the time varying pitch modulations are different between deaf and hearing males: hearing males produce calls with more downsweeps (black circle) and deaf males with more upsweeps (gray circle). The black arrows in the scatter plot point to the example calls shown in B. B. Spectrogram of two example calls from acoustic Group 4, per male bat. Note that hearing male vocalizations are short downsweep vocalizations with a low Third Spectral Quartile (black dotted lines) while the deaf male vocalizations are longer and more pitch modulated vocalizations (presence of upsweeps) with high Third Spectral Quartile (black dotted lines). The audio file of each of these examples can be found as supplemental materials (Sound_S1.wav, Sound_S2.wav, Sound_S3.wav, Sound_S4.wav, Sound_S5.wav, Sound_S6.wav, Sound_S7.wav, Sound_S8.wav).

## Discussion

In this study, we assessed the necessity of hearing for the development of a normal vocal repertoire in Egyptian fruit bats. The acoustic analysis demonstrated that the repertoire of this species consists of a large majority of innate vocalizations (>75% for males and >95% for females). Auditory input is not necessary to establish, fine-tune or maintain the neural motor programs controlling the production of these calls. However, some vocal features were not produced by males that lacked auditory input (downsweep calls with filtered higher harmonics) and atypical spectro-temporal patterns emerged in a few calls from deaf females, indicating that neural motor programs for these calls might be plastic and require an intact auditory input to properly mature. According to the source-filter theory that was first developed to explain human voice production^18,19^ and then applied to animal vocalizations^20–22^, the upsweeps and downsweeps of the fundamental frequency are controlled by the vocal source (the larynx), while the filtering of call harmonics and formants are achieved by change in the shape of the vocal tract (e.g., vocalizing with open or closed mouth). Under this theory, the results of the present acoustic analyses support the view that male Egyptian fruit-bats need to hear themselves, and/or other bats, in order to properly control both their larynx and upper vocal tract to produce the downsweep calls. The hypothesis of a fine motor control of the larynx is consistent with our recent findings demonstrating a direct monosynaptic pathway from the vocal-motor cortex to the brainstem center innervating the larynx^12^. These later findings are, so far, the strongest evidence of the existence of this direct connection in mammals, which has been suggested as a central anatomical hallmark of the neural circuitry that enables vocal production learning^23–27^.

Previous experiments, such as pups reared in social isolation, or manipulation of the acoustic environment of both young and adults, had shown effects when considering together all vocalizations in the Egyptian fruit bat repertoire^9–11^. Deafening would also affect all the vocalizations in the adult repertoire of the pale spear-nosed bat, *Phyllostomus discolor*^28^. We cannot exclude that in our experiment, deaf bats could have compensated for their lack of hearing by using somatosensation as a sensory feedback and social responses of their hearing companions as guiding cues to produce most of the vocalizations in their repertoire. Somatosensation is indeed a feedback that is used along with audition to guide vocal production in humans and songbirds alike^29–31^. Nonetheless, even after years of communal living, males could not properly produce a significant proportion of their vocal repertoire in the absence of auditory feedback. These results pave the way to targeted studies of these male Egyptian fruit bat vocalizations, with the possibility for comparing the neural mechanisms of production between learned and innate vocalizations in a mammal brain. Furthermore, the observed difference between males and females opens new opportunities to investigate how the sex of an individual might interfere with the maturation of vocal behavior in a mammalian brain that is particularly suitable for neurophysiological research.

## Supplemental information

17 supplemental figures

24 audio clips

## Supporting information

Supplemental sounds

## Acknowledgements

The authors deeply thank the undergraduate students from UC Berkeley who worked on the first manual curation step of the bats audio recordings: Yousuf Hashmi, Sun Wee Yun, Nicole Wang,

Maggie McDonough, Leo Huang, Jason Huang, Christopher Lung, Aunesha Bhowmick, Anya Cutter, Isha Yeleswarapu, Geoffrey Li, Ethan Lee, Dorian Andrei and Chirstina Yao. We also would like to thank the undergraduate students from UC Berkeley who participated in establishing the automatic algorithm for bat call detection: Shunyu Wang, Richard Ma, Jonathan Wang and Chase Adams. We thank Yartsev lab members for discussions and thoughts on the project, L. Wiegrebe for his help on the initial project design and sharing of ABR protocol & software, and OLAC staff for support with animal husbandry and care. This research was supported by the following supports to MMY: NIH (DP2-DC016163), the New York Stem Cell Foundation (NYSCF-R-NI40), the Alfred P. Sloan Foundation (FG-2017-9646), the Brain Research Foundation (BRFSG-2017-09), the Packard Fellowship (2017-66825), the Klingenstein-Simons Fellowship (2016-040745), the Pew Charitable Trust (00029645) and The McKnight Foundation (2017-042823) (to M.M.Y.). Work in the Raphael Lab was supported by the NIH (NIDCD R01-DC014832) and the R. Jamison and Betty Williams Professorship.

## Author contributions

Conceptualization: MMY, SEM, JEE

Methodology: MMY, SEM, JEE, YR, LAB, DLS

Resources: MMY, YR Data Curation: JEE

Investigation: SEM, JEE, DG, LAB

Software: JEE, TN

Formal acoustical analysis: JEE

Formal histological analysis: LAB, DLS, YR

Funding acquisition: MMY

Supervision: JEE, MMY

Writing – original draft: JEE

Writing – review & editing: JEE, MMY, LAB, TN, SEM, DG, YR, DLS

## Methods

### Subjects and housing

All experimental procedures were approved by the Institutional Animal Care and Use Committee of the University of California, Berkeley. Nine wild-caught adult female Egyptian fruit-bats (*Rousettus Aegyptiacus*) and their ten pups (five females and five males) were used for this study. Each female gave birth to one pup except for one female that gave birth twice. The progenitor female bats were imported from Israel and housed in our lab colony at the University of California Berkeley for at least six months before being included in the experiments. Female bats were isolated in soundproof booths when they were highly pregnant (individual cage size: 10x12x20 inches). This protocol has been previously used to ensure that pups would not be exposed to any vocalization including their mother’s vocalizations^10^. Indeed, an isolated mother does not vocalize even in presence of its pup^10^. Each female was maintained with its pup in their soundproof booth until the end of the deafening protocol (see below) of the pup. After the end of the injection protocol (post-natal day 56), the young bats were moved to a collective housing room and housed together with their mothers and male bats from the colony in larger cages (21 x 32 x 35.8 inches). Each cage contained one saline pup and one kanamycin pup with their respective mothers and two adult males. At sexual maturity (7 months old), the young bats were separated from their mothers and housed in single sex cages with age matched unmanipulated bats.

When reaching 3-4 years of age, all ten experimental bats were grouped together in a large cage (180x60x60 cm) and isolated from the rest of the colony in a soundproof room for two months over which their vocalizations were recorded as detailed below. In all cages, bats were maintained on a reverse day-light cycle set at 12h/12h. Recordings of vocalizations were conducted during the dark period.

### Deafening protocol

Half of the pups followed a treatment by kanamycin (powder from Alfa Aesar diluted in PBS, pH 7.4, Gibco 10010-02B) while the other half were kept as control subjects and injected with saline solution (PBS, pH 7.4, Gibco 10010-02B). Starting on post-natal day 32 (PND32) and for 21 days, the pups were subcutaneously injected twice daily (8 hrs apart). The dosage of kanamycin was set to increase stepwise over the 21 days (6 days at 300 mg/kg, 5 days at 400 mg/kg, 5 days at 500 mg/kg and 5 days at 600 mg/kg). The weight of the young bats was measured at most 4 days before a change of dosage or at least on the eve of a change of dosage to correctly adjust dosage. Control subjects were injected with the same volume of saline solution as the volume they would have been injected with if they were kanamycin subjects.

### Audiogram

Eight months after the end of the acoustic recordings, the audiogram of each experimental bat was established by measuring their auditory brain response (ABR) to tone pips of 2.5 ms (Hann window) spanning the auditory range expected for that species (1.5 - 90kHz, 25 logarithmically spaced levels of frequencies) and presented at various levels of intensity (30-110 dB SPL, 9 steps of 10dB SPL), following the same protocol as Linnenschmidt and Wiegrebe (2019)^32^.

Briefly, bats were isolated and fasted from 1h and up to 2h before inducing anesthesia by subcutaneous injection of a mix of Ketamine (22 mg/kg), DexaMedetomidine (0.09 mg/kg) and Midazolam (0.45 mg/kg). Once the bat was deeply anesthetized, it was placed laying on its stomach on a heating pad in a soundproof room, facing the loudspeaker (Scanspeak R2604/833000) at 5 cm, with an oxygen delivery tube placed right under its nose (flow rate ∼300mL/min). The animal temperature and breathing rate were monitored every 15 min during the procedure. The ABR was measured as the difference of potential between a signal subdermal electrode inserted at the base of the head in the neck of the animal and a reference subdermal electrode inserted subcutaneously atop of the head between the two ears. The whole system was grounded by an alligator clip placed on the lower part of one ear. The electrode signals were amplified by a TDT headstage (RA4Li, Tucker Davis Technologies Inc., Fl, USA) and further amplified and digitized by an audio interface (Fireface UC, RME, Haimhausen, Germany; sampling rate 192 kHz). After bandpass filtering between 100 Hz and 3 kHz and downsampling 10 times, the electrophysiological signals were stored for offline analysis. The 225 (nine amplitude levels and 25 frequency steps) tone pips were each presented 256 times at a repetition rate of 44Hz. Using previously used matlab algorithm^32^, tone pips were generated at 192kHz, digitally equalized in magnitude and phase to compensate for the loudspeaker impulse response, converted to analog using the same audio interface as the one used for processing the ABR signal, and amplified using one channel of an audio amplifier (AVR 347, Harman Kardon, Stamford, CT, USA).

For each of the 256 presentations of a tone pip, the ABR signal was extracted in two windows of 7 ms, one aligned to the tone pips (response window) and the second one randomly placed between 0 and 23ms after stimulus onset (background noise window). From these 256 pairs of snippets of signal, we generated one tone response (the average electrophysiological signal over all response windows) and one background response (the average electrophysiological signal over all background windows) and measured the RMS of each. We bootstrap the calculation of the RMS background response for each stimulus by repeating 1000 times the selection of random background noise windows in the 256 ABR time series. The exact p-value of obtaining a tone response RMS higher than the background response RMS values was then calculated as the number of background response RMS values equal or higher than the tone response RMS for that stimulus. We could then draw the audiogram for each bat as the lowest sound amplitude, for each carrier frequency tested, at which the RMS of the ABR signal was significant at p=0.05 (see Figure S14).

### Histology

#### Cochlear extraction

After ABR measurements, the bats still under anesthesia were euthanized by a lethal intraperitoneal injection of sodium pentobarbital. The body was then transcardially perfused with 200mL of phosphate buffer saline (PBS) solution (pH=7.4) spiked with 0.5mL of Heparin (1,000 USP units/mL) followed by 250 mL of fixative (3.7% formaldehyde PBS solution, pH=7.4). The temporal bones were removed from the skull and the inner ears isolated. The oval and round window were opened to facilitate flow of fixative solution for rapid fixation. Inner ear tissues were shipped to the University of Michigan for histological analyses.

The cochleae with their surrounding bone were decalcified in 5% EDTA until the bone was soft to facilitate dissection of the sensory area, the organ of Corti. The tissues were rinsed in PBS and further dissection was done to remove the bone, stria vascularis and tectorial membrane.

#### Phalloidin staining

The cochleae were permeabilized with 0.3% Triton X-100 in PBS for 15 minutes followed by incubation with Alexa Fluor 488 phalloidin (Invitrogen A12379) for 40 minutes, all at room temperature. After rinsing with PBS, the cochleae were further dissected into segments of the basal turn of the cochlea (higher frequencies area) and the apical turn (lower frequencies). Segments were then mounted on slides with Prolong Gold (Invitrogen P36930). The slides were stored in the dark at 4°C until viewed with fluorescence microscopy.

#### Imaging of the organ of Corti

The samples were imaged with a Leica DMRB epifluorescence microscope. Images were captured using a CCD cooled SPOT-RT monochrome digital camera. Digital images were processed with Adobe Photoshop.

### Audio recordings

When the animals reached 3-4 years old, the ten experimental bats were recorded for 35 days over the course of seven weeks, for 5-7 hours per day during their night cycle. For acclimation purposes, the bats were all housed in the recording cage starting 16 days before the first day of audio recording and were housed in that recording cage for the entire duration of the recording period (7 weeks). The cage was a rectangular prism (180 x 60 x 60 cm) that had two sides made of plexiglass, thereby permitting clear remote visual monitoring of bats behavior via two cameras (Flea 3 FLIR). The remaining sides of the enclosure were made of plastic mesh, allowing bats to easily perch and crawl on the surfaces. The enclosure was placed in an acoustically shielded room and all recording sessions were conducted during the dark cycle under red LED light. All bats were equipped daily with audio data-loggers to record and identify their vocalizations. Water was given *ad libitum* and the daily diet supply was placed into the cage at the beginning of each recording session to increase the diversity of behavioral contexts. The audio recordings of each individual bat vocalizations were performed using both an audio data-logger and an ambient ultrasonic microphone (Earthworks, M50) centered 20 cm above the cage ceiling and connected to the main computer unit via an analog to digital convertor (MOTU, 896mk3). The audio collected from the ambient microphone was recorded throughout the session (sampling rate of 192 kHz) using an in-house Matlab GUI (VocOperant; https://github.com/julieelie/operant_bats). As previously described (Rose et al 2021) the audio data-logger consists in a single-axis, low mass, piezo-ceramic accelerometers (BU-27135, Knowles Electronics, sensitivity 0-10kHz) that is mounted on a flexible rubber necklace placed against the throat of the subject, in a way that does not restrict normal behavior but enables the detection of laryngeal vibrations produced during vocalizations. The signal of the accelerometer is recorded, digitized at a sampling rate of 50 Hz, and saved on removable SD cards with a wireless audio data logging device (“audiologgers”; Audio Logger AL1 or AL2, Deuteron Technologies) mounted on the necklace on the back of the subject. The audiologger and its lithium-polymer battery are encapsulated in a house-made 3D-printed plastic casing to prevent damage to the electronics. All audiologgers were controlled and synchronized by a single transceiver. The Egyptian fruit bats in our experiment weighed more than 110 gr and could fly with ease while equipped with their audiologger. The synchronization between the microphone recording and the transceiver controlling the audiologgers was achieved using transistor-transistor logic (TTL) pulses generated by an UltraSoundGate Player 216H (Avisoft Bioacoustics) and sent via coaxial cables. The experimenter could monitor the behavior of the animals in an ante-chamber via the video cameras and the ambient microphone.

### Vocalizations detection and acoustic analysis

From all the days of recording, we sampled 13 days for acoustical analysis. These days were chosen such as to optimize their spread across the 8 weeks of recording and optimize the recording conditions (no audiologger breaking incident).

#### Vocalization detection

Acoustic data logged as voltage traces on the SD cards of the audiologgers were extracted into Matlab files and aligned across bats simultaneously recorded using a custom-made Matlab code (https://github.com/NeuroBatLab/LoggerDataProcessing/). Potential vocalizations were then detected and segmented from these piezo recordings using an in-house series of Matlab scripts (https://github.com/NeuroBatLab/SoundAnalysisBats). The whole process consisted in three major steps: 1) detection of sound events on audiologger signals, 2) automatic classification between vocalizations and noise, 3) manual curation of potential vocalizations. Figure S15 presents a full flow chart of the approach taken for the detection of vocalizations in audiologger recordings.

1. Detection: To focus on the detection of vocalizations emitted by the bat wearing the collar, the signal of each audiologger was first band-passed between 1 and 5kHz. As previously described, this frequency range is not contaminated by airborne vocalizations from other bats standing close to the collar wearer (Rose et al, 2021). After determining a noise threshold from sections of silence during the recording session for that audiologger (three times the RMS of silence sections), potential vocalizations were detected by threshold crossing on the amplitude envelope (RMS, sampling frequency 1kHz) and any sound event above threshold for longer than 7ms was kept.
2. Automatic step of data sorting between actual vocalizations and noise: Sound events closer than 50ms were merged as a single sound sequence and a battery of 20 acoustic measurements (pre-defined acoustic features, PAFs) were applied on them (see Elie and Theunissen, 2016, for mathematical definitions): RMS, maximum amplitude, the five first momentum of the amplitude envelope taken as a distribution (mean, standard deviation, kurtosis, skewness, and entropy), the five first momentum of the frequency spectrum taken as a distribution, the three quartiles of the frequency spectrum, the mean pitch saliency, and four parameters that pertain to the sound as recorded from the ambient microphone (RMS, mean and maximum of the amplitude envelope, and maximum value of cross-correlation between the microphone and the audiologger signals). Potential vocalizations among the detected sound events were then identified using these 20 acoustic parameters as input to a support vector machine (SVM) trained on the data of two sessions manually sorted between vocalizations and noise by an expert (JEE). This data was obtained from a total of 100 min of recording of 8 bats (800 min of audiologger data) and contained 24,633 sound elements, of which 1,377 were vocalizations. The automatic sorting was set to be very conservative of vocalizations by setting the threshold on the posterior probability of a vocalization at 0.002, with the drawback of yielding many false discoveries (percentage of vocalizations correctly identified in 10-fold cross validation: 98.3%; percentage of noise miss-classified as vocalizations in 10-fold cross validation: 7.5%). After the manual curation (see below) of a reasonable dataset (five days), an extra step of automatic detection could be added to the pipeline to further filter out sequences devoid of vocalization. We trained a neural network on the manual annotations obtained from these five days of recording (4,936 vocalizations). The network was implemented using the PyTorch library and the architecture was composed of ResNet-50 with a sigmoid output layer. For each 100ms non-overlapping window of the audiologger signal, we bandpass filtered the signal between 100Hz and 5kHz, and calculated its spectrogram using *soundsig* from the BioSound library (https://github.com/theunissenlab/soundsig) with the following parameters: sampling rate=1kHz; frequency spacing=100Hz; dynamic range=50dB. The spectrogram was then colorized using the color scale in *soundsig*. The training label for each 100ms window was 1 if there existed a vocalization in that window and 0 otherwise. The 3-channel color spectrograms and corresponding binary labels were used as inputs for the network. To take into account the large imbalance between noise and vocalization elements in the dataset, each training batch was set to contain 5 noise elements for each vocalization element. We used a weighted binary cross-entropy loss function with penalty=7.5 to further address the imbalance within each training batch. 20 percent of the data was held out for validation. We randomly initialized the network weights and trained the network for 10 epochs using the Adam optimizer with learning rate=1e-3. A sound sequence on a given audiologger was considered as worthy of manual examination if the maximum of all posterior probability values (one value per 100ms window) was higher than 0.9. At this threshold of maximum posterior probability value, the trained neural network achieved high performance of correct classification of sequences containing vocalizations (98.6%) with low values of false discovery (5.4% of noise sequences detected as containing vocalizations) enabling to greatly speed up the vocalization discovery procedure. The data from 8 more days of recording were evaluated by the trained neural network.
3. Manual curation: to further eliminate noise and check the identity of the bat producing vocalizations, each potential vocalization was aurally and visually scrutinized by trained students and an expert (JEE) based on the inspection of the spectrograms of its signal as recorded from the ambient microphone and from the audiologgers of all the bats (*manual annotation* in Figure S15). Vocal utterances had to have a minimum duration of 7ms to be considered as vocalizations and those spaced by less than 50ms were grouped together. A final step of audio quality evaluation performed by an expert eliminated vocalizations that were contaminated by other vocalizations or noise on the microphone recording.

#### Acoustic features and modulation power spectrum (MPS)

28 predefined acoustic features (PAFs) were chosen to describe both the temporal and the spectral content of vocalizations. For each vocalization, PAFs were calculated from both the audiologger recording and from the ambient microphone recording. The signal captured by the piezo sensor of the audiologger was particularly suitable to measure pitch or amplitude related acoustic features because it is consistently placed very close to the larynx of the animal, the vocal source. The PAFs calculated from the audiologger signal were: duration, pitch saliency (a measure of how tonal vs noisy a sound is), fundamental frequency, coefficient of variation (CV) of the fundamental frequency, the five first momentum of the sound temporal envelope taken as a distribution (mean, standard deviation, kurtosis, skewness and entropy), the amplitude periodicity frequency (frequency peak in the amplitude power spectrum), the amplitude periodicity power index (ratio of the power at the amplitude periodicity frequency with the signal RMS), the five first momentum of the sound power spectrum taken as a distribution (mean, standard deviation, kurtosis, skewness and entropy) and the first, second and third spectral quartiles. The PAFs calculated from the microphone signal were: the five first momentum of the sound spectrum taken as a distribution (mean, standard deviation, kurtosis, skewness and entropy) and the first, second and third spectral quartiles. All these parameters except the two amplitude periodicity measurements were calculated using *soundsig* from the BioSound Python library (https://github.com/theunissenlab/soundsig). The two amplitude periodicity measurements were calculated using custom Matlab code.

The modulation power spectrum (MPS) of each vocalization was calculated from the microphone signal using *soundsig* from the BioSound Python library (https://github.com/theunissenlab/soundsig). The MPS is the 2D fast Fourier transform of the sound spectrogram. To ensure that each vocalization MPS could be calculated at the same time resolution, we ensured that each vocalization was isolated within a 100 ms window. After bandpassing the microphone signal between 0.3 kHz and 50 kHz, the spectrogram was calculated at a 1 kHz sampling rate using a frequency spacing of 100Hz. The spectrogram was then z-scored and the MPS calculated using a window bin of 100ms. The MPS range was focused on values with the most energy: the range of spectro-temporal modulations was restricted from 0 to 3 cycles/kHz for the frequency modulations and from -150 Hz to 150 Hz for the temporal modulations (see example MPS in Figure S9). To account for the various dynamic range across the spectro-temporal modulations, the MPS of each vocalization was normalized by dividing with the average MPS over all vocalizations in the dataset to obtain the Normalized Modulation Power Spectrum (NMPS, Figure S9).

#### UMAP projection and acoustic clustering

The NMPS of the 28,091 vocalizations were reduced to fewer dimensions by applying the algorithm UMAP (Uniform manifold approximation and projection) setting the number of neighbors to 10, the minimum distance to 0.051 and the target weight to 0.55 (Matlab code provided by the Herzenberg Lab at Stanford University, CA, USA). The NMPS as represented by the UMAP values were then used as input to a hierarchical agglomerative clustering (HAC) algorithm (*linkage* function of Matlab using the Ward method and the Euclidean metric) to divide the acoustic space covered by the vocalizations into groups. The optimal number of dimensions for the UMAP, the optimal tree cut and optimal number of acoustic groups were found by measuring the average and median silhouette value over UMAPs of various dimensions and various levels of tree cutting in the HAC (Figure S10). Four acoustic groups in a 3D UMAP were found to be optimal numbers.

#### Vocal dispersion

Vocal dispersion is calculated for each individual bat as the generalized variance of its vocalizations using the first 100 dimensions of the PCA on the NMPS of vocalizations from the whole dataset. The first 100 dimensions explained 90% of the variance. The result depicted on Figure S5A, showing a larger vocal dispersion for males as compared to females, was robust across all the number of dimensions tested (from 10 to 100 with a step of 10 and from 100 to 300 with a step of 20).

#### Acoustic dissimilarity (distance to opposite sex and distance to opposite hearing ability)

The dissimilarity was calculated for each vocalization emitted by bat *i* as the distance to the vocalizations from the nearest target bats *j* (*j* ≠ *i)* in the space of the principal component analysis (PCA) calculated on NMPS of vocalizations (Figure 2). Dimensions of the PCA had to explain at least 1% of the variance to be included in this measure to ensure that the measure would not be noise sensitive. As such the first 5 dimensions were retained. To represent how males and females differentially occupy the acoustic space (Figure 2A), the target bats were the opposite sex bats. To represent which vocalizations were affected by deafening (Figure 2B and 2D), the target bats were the opposite treatment but same sex bats. For each vocalization, the acoustic dissimilarity is the distance to the closest of the target bats beyond what would be expected given the local density of bats vocalizations. The distance from one particular vocalization of bat *i* to the vocalizations of a *j* target bat was calculated as the average Euclidean distance to the nearest 10 vocalizations of target bat *j*. The local density of bat vocalizations (null distance) was the distance to any bat for the calculation of the sex dissimilarity, and the distance to any same-sex bat for the calculation of hearing/deaf dissimilarity. For a vocalization emitted by a male, the sex dissimilarity is the difference between the average Euclidean distance to the 10 nearest vocalizations from the closest female bat, with the average Euclidean distance to the 10 nearest vocalizations from any closest bat. For a vocalization emitted by a hearing male, the hearing/deaf dissimilarity is the difference between the average Euclidean distance to the 10 nearest vocalizations from the closest deaf male bat, with the average Euclidean distance to the 10 nearest vocalizations from the closest male bat (hearing or deaf). The proportion of the repertoire that was affected by deafening was estimated as the proportion of vocalizations that displayed a hearing/deaf dissimilarity value larger than the null distance to any same sex bats (Figure S7).

#### Regularized Permutation Discriminant Function Analysis

The performance of classification of NMPS of vocalizations according to the hearing ability of the emitter bat (Deaf and Hearing) was determined using a regularized permutation discriminant function analysis (RPDFA). For each comparison (set of data), this procedure consisted in a regularization achieved by a principal component analysis (PCA) on the NMPS, followed by a Linear Fisher Discriminant Function Analysis (DFA). The performance of the DFA was estimated in 10-fold cross-validation fixing the prior probability at 0.5. To find out the optimal number of PCA dimensions to consider in the DFA, the performance of the classifier was measured across a range of numbers of dimensions up to the number of dimensions that explained 90% of the variance in the PCA (step of 10 up to 100 dimensions; step of 20 between 100 and 500 dimensions; step of 100 dimensions above). The optimal number of dimensions of the PCA to consider in the DFA was found as the first value where the increase in classification performance weighted by the increase of percentage of variance explained by the change of number of dimensions is smaller than half the average standard deviation of the performance across all values of dimensions tested (see Figures S4, S6, S16 and S17). The performance of the classifier at this optimized number of PCA dimensions was retained and reported in the article. The chance level for the classification performance using the optimized number of PCA dimensions was determined by running a permutation procedure where the treatment of individuals was randomized across bats. Depending on the number of subjects in the dataset there were 126 (ten male and female bats), 9 (six female bats) or 2 (four male bats) different possible permutations. To determine if the odds of correct classification was higher in the non-permuted treatment case than in the permutation cases, we perform a right tailed Fisher’s exact test for each permutation.

For display and interpretation purposes, the DFA at the optimal number of PCA dimensions was recalculated without cross-validation to obtain the discriminant function as shown in Figure 3 and Figure S11.

## References

1. Mahrt, E.J., Perkel, D.J., Tong, L., Rubel, E.W., and Portfors, C.V. (2013). Engineered Deafness Reveals That Mouse Courtship Vocalizations Do Not Require Auditory Experience. Journal of Neuroscience 33, 5573–5583. 10.1523/JNEUROSCI.5054-12.2013.

2. Tyack, P.L. (2020). A taxonomy for vocal learning. Philosophical Transactions of the Royal Society B: Biological Sciences 375, 20180406. 10.1098/rstb.2018.0406.

3. Tyack, P.L. (2016). Vocal Learning and Auditory-Vocal Feedback. In Vertebrate Sound Production and Acoustic Communication, R. A. Suthers, W. T. Fitch, R. R. Fay, and A. N. Popper, eds. (Springer International Publishing), pp. 261–295. 10.1007/978-3-319-27721-9_9.

4. Janik, V.M., and Slater, P.J.B. (2000). The different roles of social learning in vocal communication. Animal Behaviour 60, 1–11. 10.1006/anbe.2000.1410.

5. Fischer, J., Wegdell, F., Trede, F., Dal Pesco, F., and Hammerschmidt, K. (2020). Vocal convergence in a multi-level primate society: insights into the evolution of vocal learning. Proceedings of the Royal Society B: Biological Sciences 287, 20202531. 10.1098/rspb.2020.2531.

6. Ruch, H., Zürcher, Y., and Burkart, J.M. (2018). The function and mechanism of vocal accommodation in humans and other primates. Biological Reviews 93, 996–1013. 10.1111/brv.12382.

7. Wirthlin, M., Chang, E.F., Knörnschild, M., Krubitzer, L.A., Mello, C.V., Miller, C.T., Pfenning, A.R., Vernes, S.C., Tchernichovski, O., and Yartsev, M.M. (2019). A Modular Approach to Vocal Learning: Disentangling the Diversity of a Complex Behavioral Trait. Neuron 104, 87–99. 10.1016/j.neuron.2019.09.036.

8. Janik, V.M., and Knörnschild, M. (2021). Vocal production learning in mammals revisited. Philosophical Transactions of the Royal Society B: Biological Sciences 376, 20200244. 10.1098/rstb.2020.0244.

9. Prat, Y., Azoulay, L., Dor, R., and Yovel, Y. (2017). Crowd vocal learning induces vocal dialects in bats: Playback of conspecifics shapes fundamental frequency usage by pups. PLOS Biology 15, 1–14. 10.1371/journal.pbio.2002556.

10. Prat, Y., Taub, M., and Yovel, Y. (2015). Vocal learning in a social mammal: Demonstrated by isolation and playback experiments in bats. Science Advances 1, e1500019. 10.1126/sciadv.1500019.

11. Genzel, D., Desai, J., Paras, E., and Yartsev, M.M. (2019). Long-term and persistent vocal plasticity in adult bats. Nature Communications 10, 3372. 10.1038/s41467-019-11350-2.

12. Wirthlin, M.E., Schmid, T.A., Elie, J.E., Zhang, X., Shvareva, V.A., Rakuljic, A., Ji, M.B., Bhat, N.S., Kaplow, I.M., Schäffer, D.E., et al. (2022). Vocal learning-associated convergent evolution in mammalian proteins and regulatory elements (Neuroscience) 10.1101/2022.12.17.520895.

13. Rose, M.C., Styr, B., Schmid, T.A., Elie, J.E., and Yartsev, M.M. (2021). Cortical representation of group social communication in bats. Science 374, eaba9584. 10.1126/science.aba9584.

14. Yartsev, M.M., and Ulanovsky, N. (2013). Representation of Three-Dimensional Space in the Hippocampus of Flying Bats. Science 340, 367–372. 10.1126/science.1235338.

15. Liberti, W.A., Schmid, T.A., Forli, A., Snyder, M., and Yartsev, M.M. (2022). A stable hippocampal code in freely flying bats. Nature 604, 98–103. 10.1038/s41586-022-04560-0.

16. Nottebohm, F. (1972). The Origins of Vocal Learning. The American Naturalist 106, 116– 140.

17. Singh, N.C., and Theunissen, F.E. (2003). Modulation spectra of natural sounds and ethological theories of auditory processing. The Journal of the Acoustical Society of America 114, 3394–3411. 10.1121/1.1624067.

18. Fant, G. (1980). The Relations between Area Functions and the Acoustic Signal. Phonetica 37, 55–86. doi:10.1159/000259983.

19. Fant, Gunnar (1971). Acoustic theory of speech production: with calculations based on X-ray studies of Russian articulations.

20. Vannoni, E., and McElligott, A.G. (2007). Individual Acoustic Variation in Fallow Deer (Dama dama) Common and Harsh Groans: A Source-Filter Theory Perspective. Ethology 113, 223–234. 10.1111/j.1439-0310.2006.01323.x.

21. Beckers, G.J.L., Nelson, B.S., and Suthers, R.A. (2004). Vocal-Tract Filtering by Lingual Articulation in a Parrot. Current Biology 14, 1592–1597. 10.1016/j.cub.2004.08.057.

22. Taylor, A.M., and Reby, D. (2010). The contribution of source–filter theory to mammal vocal communication research. Journal of Zoology 280, 221–236. 10.1111/j.1469-7998.2009.00661.x.

23. Jarvis, E.D. (2019). Evolution of vocal learning and spoken language. Science 366, 50–54. 10.1126/science.aax0287.

24. Simonyan, K. (2014). The laryngeal motor cortex: its organization and connectivity. Current Opinion in Neurobiology 28, 15–21. 10.1016/j.conb.2014.05.006.

25. Jürgens, U. (2002). Neural pathways underlying vocal control. Neuroscience & Biobehavioral Reviews 26, 235–258. 10.1016/S0149-7634(01)00068-9.

26. Fitch, W.T. (2018). The Biology and Evolution of Speech: A Comparative Analysis. Annu. Rev. Linguist. 4, 255–279. 10.1146/annurev-linguistics-011817-045748.

27. Kuypers, H.G.J.M. (1958). Corticobulbar connexions to the Pons and lower brain-stem in man: an anatomical study. Brain 81, 364–388. 10.1093/brain/81.3.364.

28. Lattenkamp, E.Z., Linnenschmidt, M., Mardus, E., Vernes, S.C., Wiegrebe, L., and Schutte, M. (2021). The vocal development of the pale spear-nosed bat is dependent on auditory feedback. Philosophical Transactions of the Royal Society B: Biological Sciences 376, 20200253. 10.1098/rstb.2020.0253.

29. Nasir, S.M., and Ostry, D.J. (2008). Speech motor learning in profoundly deaf adults. Nature Neuroscience 11, 1217–1222. 10.1038/nn.2193.

30. Tremblay, S., Shiller, D.M., and Ostry, D.J. (2003). Somatosensory basis of speech production. Nature 423, 866–869. 10.1038/nature01710.

31. McGregor, J.N., Grassler, A.L., Jaffe, P.I., Jacob, A.L., Brainard, M.S., and Sober, S.J. (2022). Shared mechanisms of auditory and non-auditory vocal learning in the songbird brain. eLife 11, e75691. 10.7554/eLife.75691.

32. Linnenschmidt, M., and Wiegrebe, L. (2019). Ontogeny of auditory brainstem responses in the bat, Phyllostomus discolor. Hearing Research 373, 85–95. 10.1016/j.heares.2018.12.010.

