## Supplemental figures for "Role of auditory feedback for vocal production learning in the Egyptian fruit-bat"

**This PDF file includes:**

Figures S1 to S17

**Other supporting materials for this manuscript include the following:**

Sounds S1 to S24

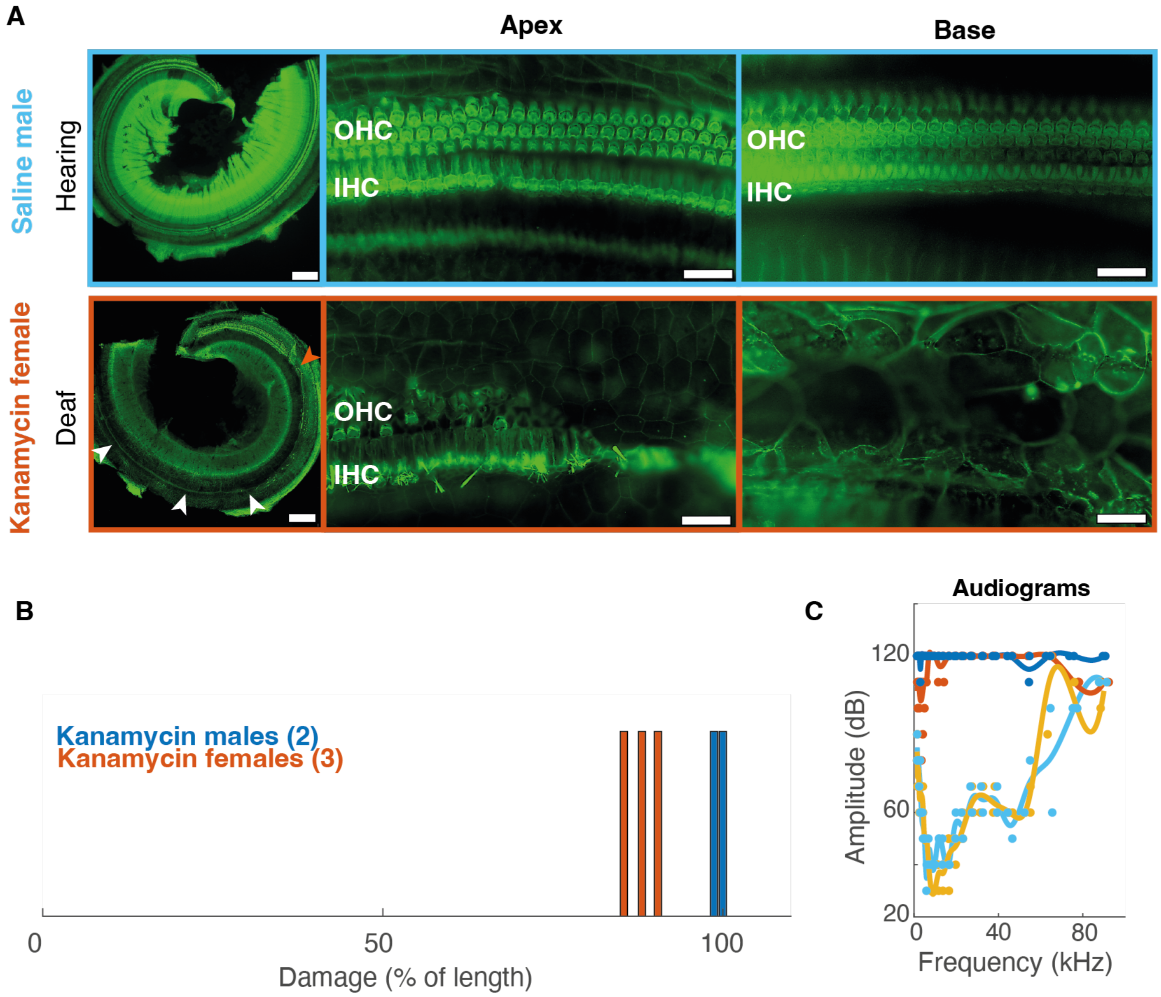


**Fig. S1.** Histological damage of kanamycin on cochlea.

A. Representative example pictures of the cochlear sensory epithelium of two bats stained with phalloidin (first row, saline injected male bat; second row, kanamycin injected female bat). The first column shows the apical turn of the organ of Corti (scale bar is 200µm). The second and third columns are higher magnification images taken from the apical and basal turns of the cochlea (scale bar is 25µm). Note in the saline bat's organ of Corti, the epithelium is intact, with three rows of outer hair cells (OHC) and one row of inner hair cells (IHC) both in the apical and basal turns. In contrast, in the kanamycin bat's cochlea, hair cells are replaced by a non-sensory flat epithelium throughout the organ of Corti (see white arrows) except in the extreme apical end of the apical turn in female kanamycin treated bats. The orange arrow points at the transition zone between flat epithelium and the apical tip where few rudimentary hair cells survived.
B. Measured tissue damage as a percentage of the cochlea length where the organ of Corti is replaced by a flat epithelium in kanamycin treated bats (values are average over the two ears). Note that at least 84% of the cochlea length was deprived of hair cells in both ears of these animals. Males were completely deaf with more than 98% of damaged tissue in both ears, while females had some remaining healthy hair cells at the apex of their cochlea, which is a region sensitive to low frequency sounds.
C. Audiograms obtained from measuring the Auditory Brainstem Response of all bats (n=10) to tone pips covering the species auditory range. Individual dots depict individual values, while lines show the average per sex and treatment fitted with a spline. Note that responses of the auditory nerve were absent or only detectable for very loud sounds (>90dB) in female kanamycin treated bats, while the expected audiogram was obtained for saline treated bats.

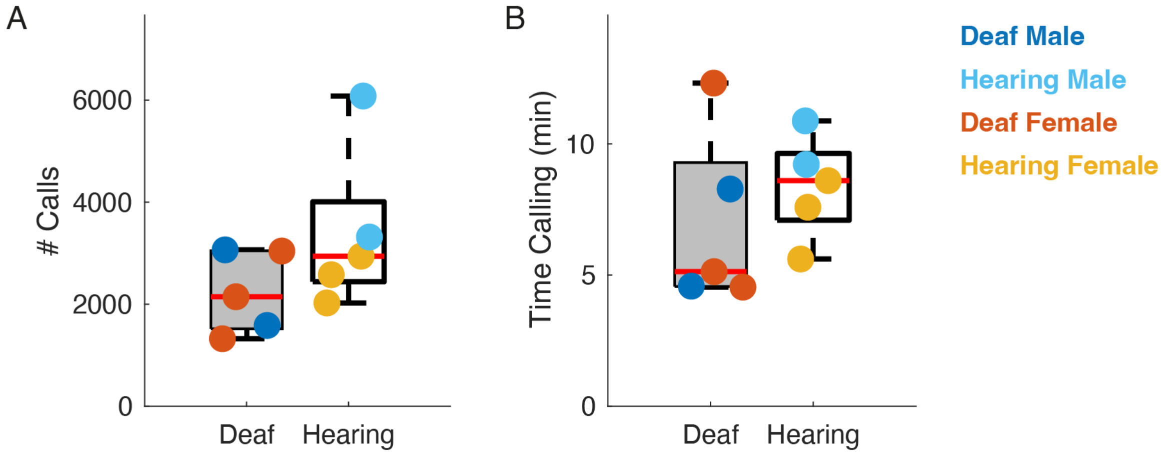


**Fig. S2.** Bat vocal activity over 13 days.

A. Box plot of the total number of vocalizations produced by bats over the 13 days of audio recording analyzed.
B. Box plot of the total duration of vocal activity of bats over the 13 days of audio recording analyzed.
Note that the dataset contains a minimum of 1,000 calls per bat or 4.5 min of vocalizing behavior. Values for each bat are depicted by the dots, color coded according to sex and hearing ability. Hearing ability had no effect on the number of calls or the time spent calling (two-sided Wilcoxon rank sum test, $p = 0.3095$ for both).

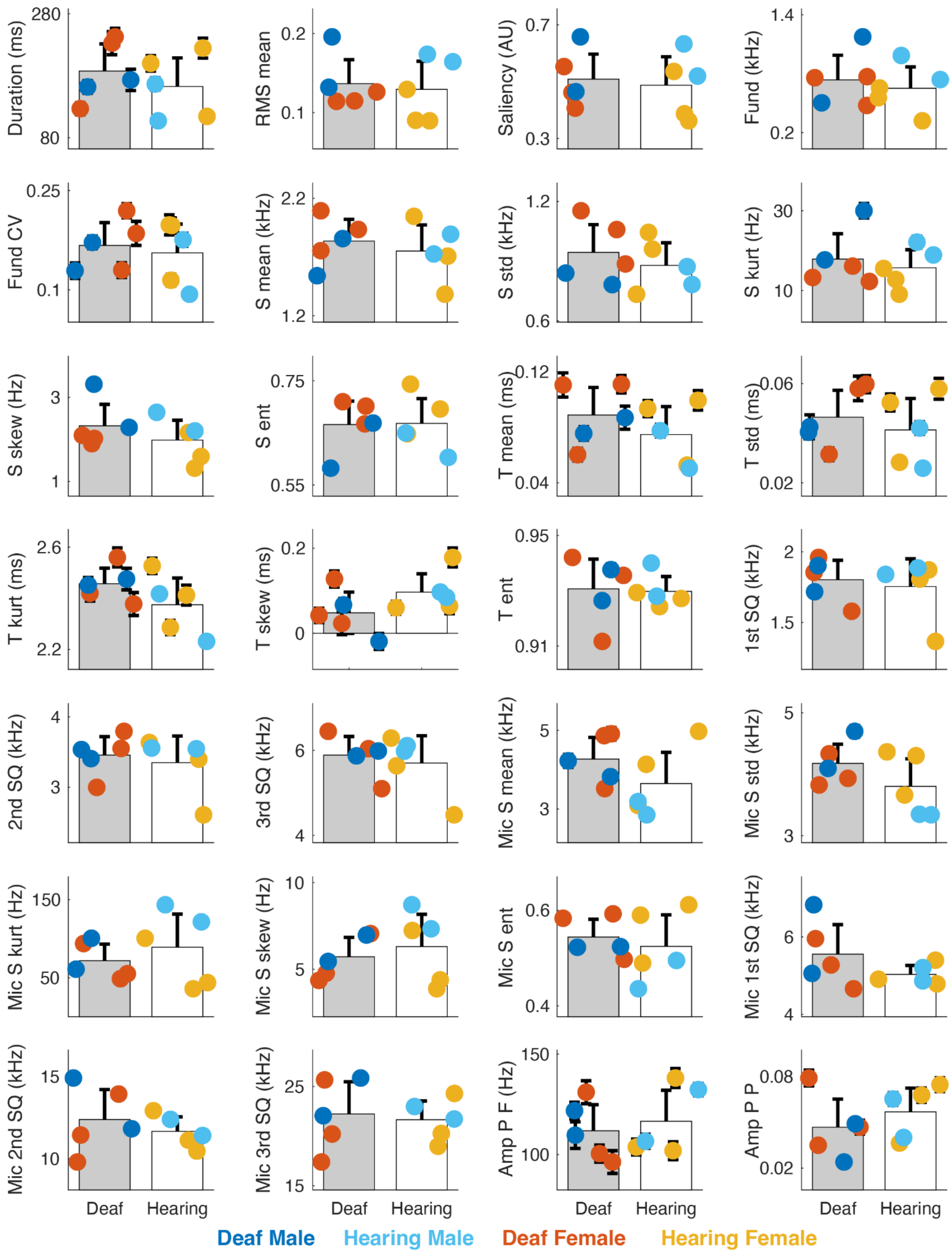


**Fig. S3.** Average acoustic features of bat calls across their vocal repertoire.

28 acoustic features were chosen to describe both the temporal and the spectral content of vocalizations (methods). Average values for each bat are depicted by the dots, color coded according to sex and hearing ability. The bars depict the average values across deaf (gray) and hearing (white) bats. Error bars are ± 2*SE. None of the acoustic parameters was significantly

different between deaf and hearing bats in a log likelihood ratio test between the full Linear Mixed Effect model with hearing ability as a predictor and bat identity as a random variable, and the reduced model missing the fixed effect hearing ability (all $p>0.05$). Acoustic features: Duration; RMS mean, average RMS of the call; Saliency, measure of pitch saliency; Fund, Fundamental frequency; Fund CV, coefficient of variation of the fundamental frequency; S mean, Spectral mean; S std, Spectral standard deviation; S kurt, Spectral kurtosis; S skew, Spectral skewness; S ent, Spectral entropy; T mean, temporal mean; T std, temporal standard deviation; T kurt, temporal kurtosis; T skew, temporal skewness; T ent, temporal entropy; 1st SQ, first spectral quartile; 2nd SQ, second spectral quartile; 3rd SQ, third spectral quartile; Amp P F, frequency of the amplitude periodicity; Amp P P, power of the amplitude periodicity.

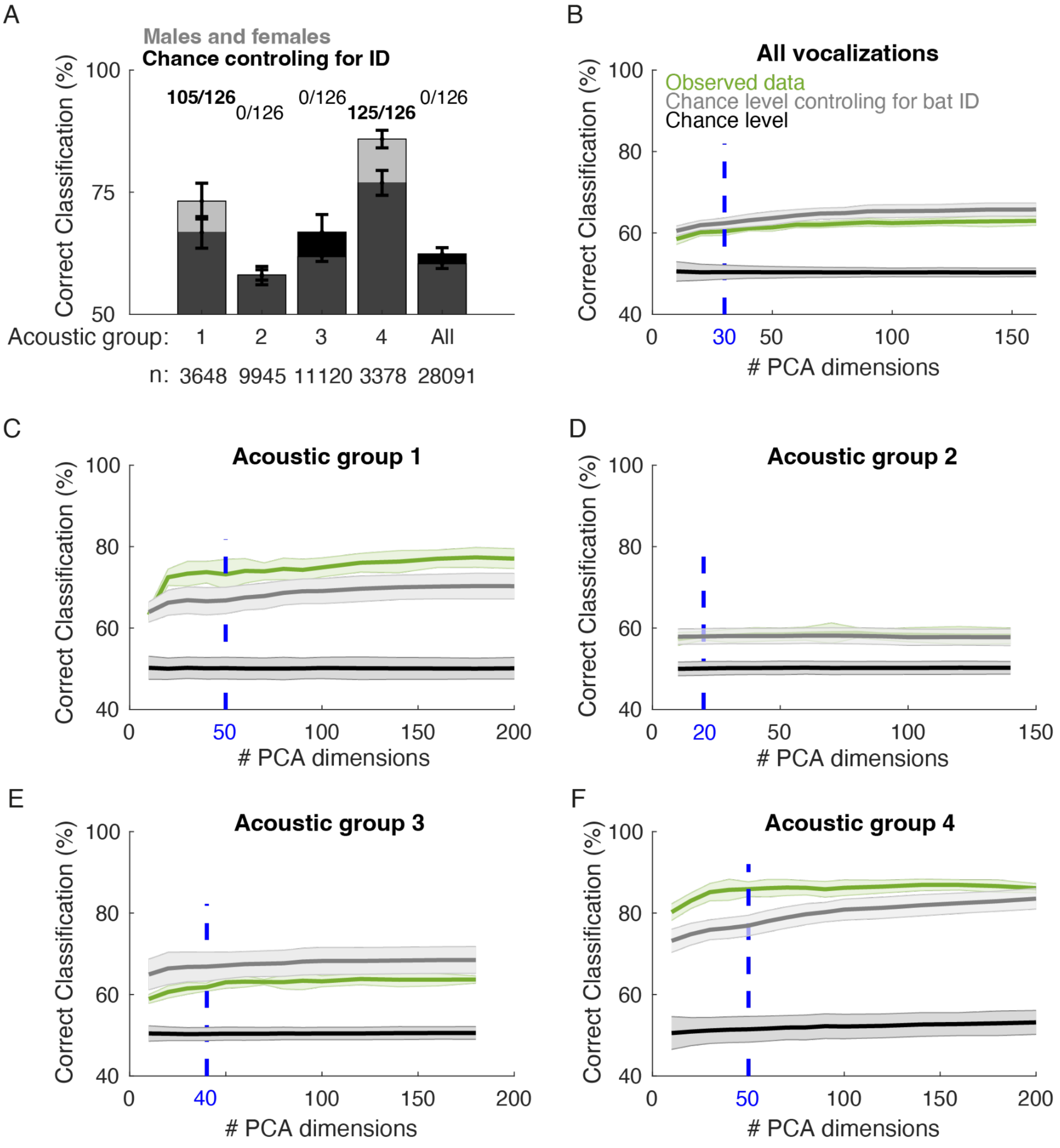


**Fig. S4.** Performance of the regularized linear classifier (RPDFA) at discriminating vocalizations produced by deaf bats from those produced by hearing bats.

A. Percentage of correct classification of vocalizations according to hearing ability (deaf or hearing) obtained for all vocalizations in the dataset ("All") and for each acoustic group (see methods and Figure 2C). Vocalizations emitted by males and females are considered together. The classification performance in permutation tests taking into account the identity of bats are shown as black bars. The bar height shows the mean performance, and error bars depict the standard deviation over 10 cross validation loops and, in the case of the permutation tests, over all 126 possible permutations of bats' hearing ability (random groups of five bats) and their 10 cross validation loops. The values above each bar indicate the number of significant exact right tailed Fisher's tests comparing the classification performance obtained with the true hearing labels (Hearing vs Deaf) and every possible permutation of these labels across individuals (126 different permutations). Note that vocalizations of deaf bats could only be discriminated from

those of hearing bats in Acoustic Group 1 and 4, but not when considering all Acoustic Groups together (‘All’).
B-F. Robustness of the classifier with respect to the dimensionality reduction step, for all vocalizations and per acoustic group. The percentage of correct classification of vocalizations between deaf and hearing bats obtained in cross-validation (10 folds) is depicted as a function of the number of PCA dimensions used to represent the MPS of vocalizations (green lines mean ± std). The optimal regularization (optimal number of PCA dimensions; dotted blue line) is chosen as a compromise between the increase of classification performance and the decrease in the percentage of variance explained in the PCA by the added new dimensions. The performance of the classifier is compared against two permutation tests: a permutation test of the hearing ability label at the level of vocalization irrespective of bat identity (dark gray lines mean ± std over 10 cross validation folds of 10 permutations), a permutation test of the hearing ability label at the level of the bat identity (light gray lines mean ± std over 10 cross validation folds of 126 permutations). This last permutation test gives a chance level that controls for the bat voice characteristics that enables the classifier to discriminate between any groups of two bats.

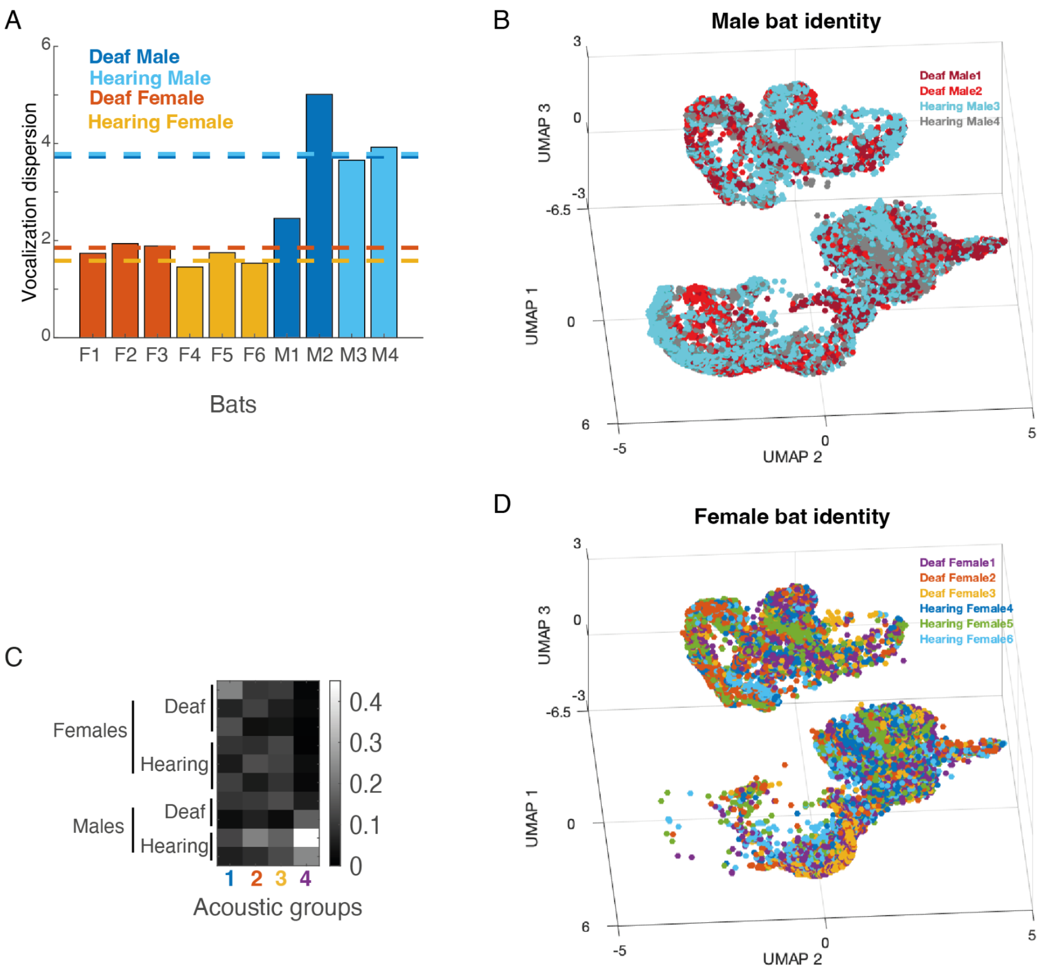


**Fig. S5.** Occupancy of the vocal space by male and female bats.
A. Vocal dispersion is the generalized variance for each bat in the PCA space applied to all vocalizations MPS in the dataset. The dotted lines give the average value across bats within each hearing ability and sex category. On average the acoustic space covered by hearing males is almost twice as large as that covered by hearing females.
B. and D. Same projection as Figure 2A with vocalizations (points) color coded by the bat identity for male bats (B) and female bats (D). Note that the vocal space with UMAP2 values below -2 is almost exclusively occupied by male calls.
C. Probability of calling within each acoustic group for each bat (the matrix sum to 1). Note that female bats rarely produced calls falling in acoustic group 4.

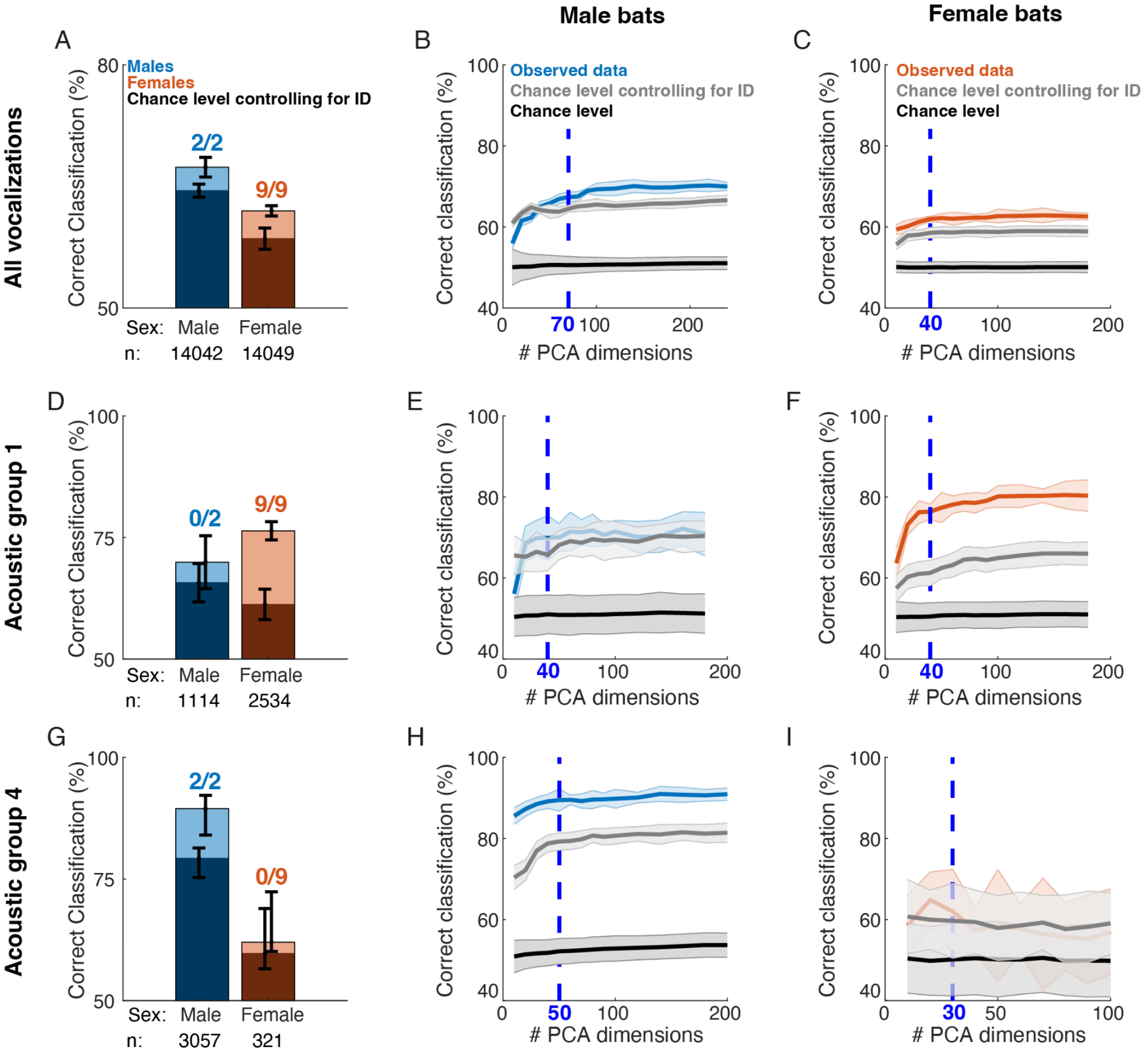


**Fig. S6.** Regularized linear classifier performance of discrimination between vocalizations from deaf and hearing bats for each sex.

A. Percentage of correct classification of vocalizations according to hearing ability (deaf or hearing) for each sex (male, blue; female, red) across all vocalizations. The classification performance of permutation tests taking into account the identity of bats are shown as black bars. The bar height shows the mean performance and error bars depict the standard deviation over 10 cross validation loops and, in the case of the permutation tests, over both 10 cross validation loops and the permutations of bats hearing ability. The number of significant ($p<0.001$) exact right tailed Fisher's tests comparing the classification performance obtained with the true hearing ability label (hearing vs deaf) and every possible permutation of these labels across individuals (2 different permutations for males, 9 for females) is shown above each bar.

B-C. Robustness of the classifiers of male calls (B) and female calls (C) with respect to the dimensionality reduction step. The percentage of correct classification of vocalizations between deaf and hearing bats obtained in cross-validation (10 folds) is depicted as a function of the number of PCA dimensions used to represent the MPS of vocalizations ("observed data", mean ± std). The optimal regularization (optimal number of PCA dimensions; dotted blue line) is chosen as a compromise between the increase of classification performance and the decrease in the percentage of variance explained in the PCA by the added new dimensions. The performance of the classifier is compared against two permutation tests: a permutation test of the hearing ability label at the level of vocalization irrespective of bat identity (dark gray lines mean ± std over 10 cross validation folds of 10 permutations), a permutation test of the hearing ability label at the

level of the bat identity (light gray lines mean ± std over 10 cross validation folds of two permutations for male calls and nine permutations for females calls). This last permutation test gives a chance level that controls for the bat voice characteristics that enables the classifier to discriminate between any groups of two bats.
D,E,F. Same as (A,B,C) for vocalizations from Acoustic Group 1.
G,H,I. Same as (A,B,C) for vocalizations from Acoustic Group 4.
The classifiers are considered to be performing above chance level when there is no overlap in error bars, all possible Fisher's tests are significant and the results are robust across the number of dimensions chosen in the regularization step. The classifiers' performance demonstrates that overall male calls and female calls are significantly affected by deafening, but only female calls are significantly affected by deafening in Acoustic group 1 and only male calls are significantly affected by deafening in Acoustic Group 4.

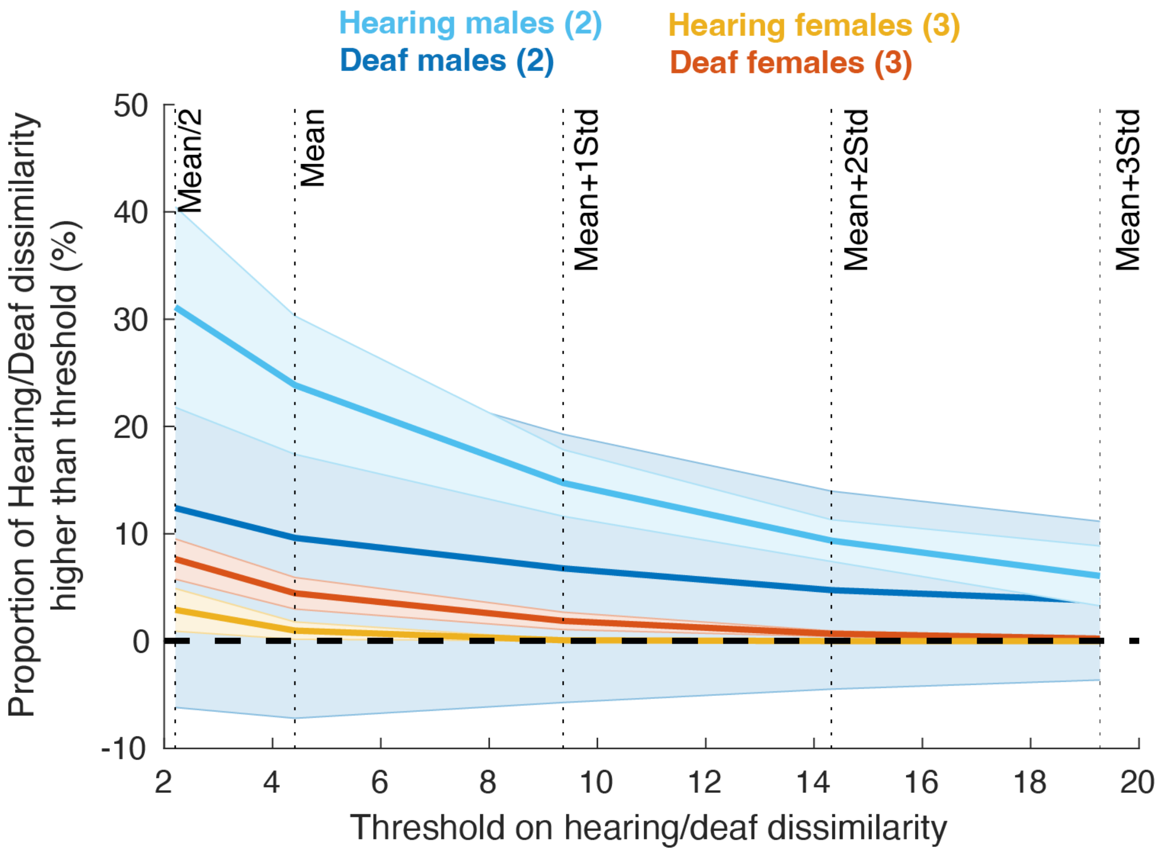


**Fig. S7.** Proportion of the repertoire with high values of hearing/deaf dissimilarity.
The average proportion of vocalizations with a hearing/deaf dissimilarity value higher than tested threshold is depicted for each sex and hearing ability (mean±ste). The tested thresholds were chosen from the statistics (mean and std) of the distribution of null distance (Euclidean distance in the 5 dimensional PCA space to any same sex bats, see methods). Note that irrespective of threshold, hearing males always have a significant proportion of their repertoire (7-30%) with high hearing/deaf dissimilarity while deaf males or hearing females proportions are never different from zero. The case of deaf females is intermediary with a small proportion of their repertoire (<10%) with high hearing/deaf dissimilarity values for lower thresholds.

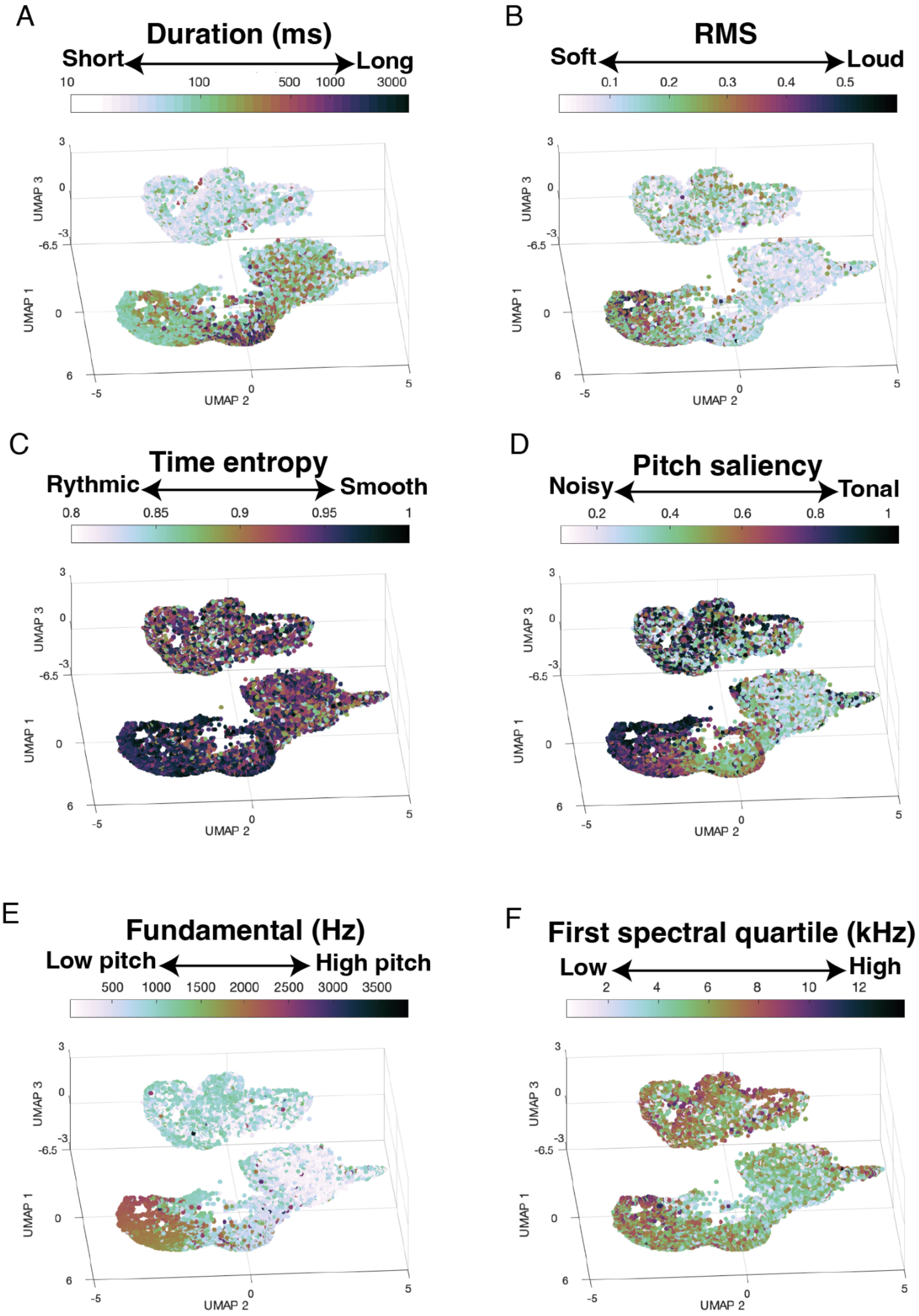


**Fig. S8.** Acoustic features variations in the vocal space covered by hearing and deaf bat vocalizations.
The Normalized Modulation Power Spectrum of each vocalization in the dataset (n=28,091) is depicted the 3-dimensional UMAP and color coded according to six acoustic features, some

calculated using BioSound (methods): the duration (A), the average amplitude as measured by the RMS (B); a measure of amplitude rhythmicity, the time entropy (C); a measure of spectral roughness, the pitch saliency (D); the average fundamental of the call (E) and the first spectral quartile (F). Note that vocalizations form two clusters, one grouping most of the short calls (<100ms) of average 1kHz fundamental frequency and another cluster grouping all the longer calls (>100ms). This later cluster group vocalizations continuously change in the vocal space from loud, smooth, highly tonal, high pitch and high spectral quartile calls (lowest values along UMAP2) to quieter, noisier, more rhythmic, lower pitch and lower spectral quartile calls (highest values along UMAP2).

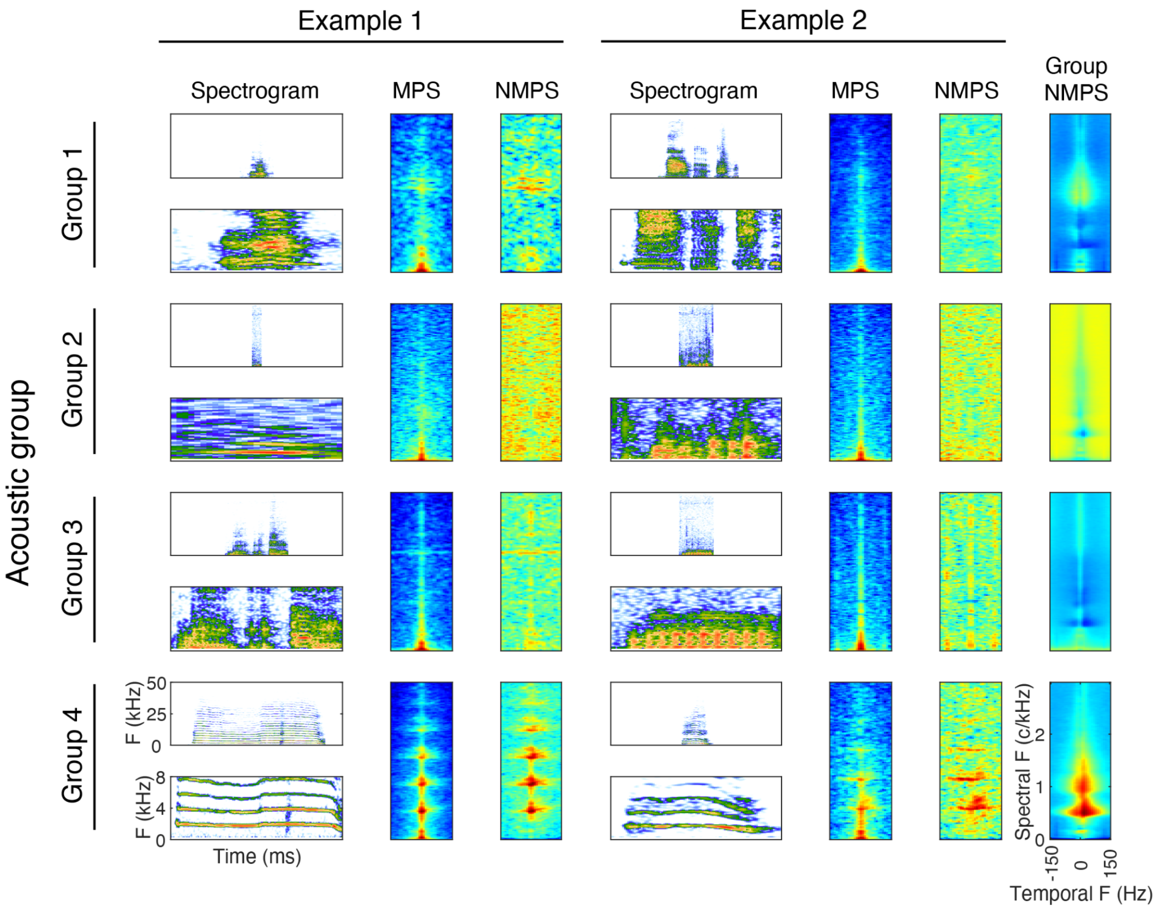


**Fig. S9.** Example vocalizations from hearing bats in each acoustic group.
For each acoustic group (see main text, methods and Figure 2C for the division of the vocalizations in acoustic groups), the spectrograms and modulation power spectra (MPS) of two example calls are given in columns. For each example call, the spectrogram used to calculate the MPS is placed in a 500ms time window and shown on the top (frequency range 0-50kHz, dB range 60), and a magnification of the spectrogram between 0 and 8kHz is presented at the bottom. The normalized modulation power spectrum (NMPS) used as input to the UMAP projection is the ratio between the MPS of each call and the average MPS over all calls in the dataset. The last column depicts the average NMPS over all calls in each Acoustic Group. The audio file of each of these examples can be found as supplementary sounds (from left to right and top to bottom: Sound S9-S16).

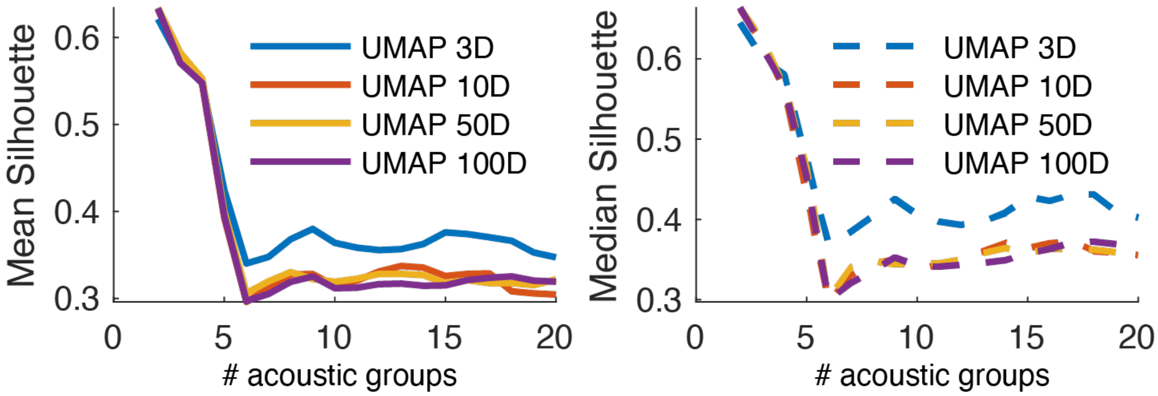


**Fig. S10.** Optimization of the procedure for parsing the acoustic space covered by vocalizations in subgroups. Four different UMAP on the NMPS of vocalizations were tested with an increasing number of dimensions (3, 10, 50 and 100 dimensions) and an increasing number of groups in the HAC (from 2 to 20 with a step of 1). The optimal number of dimensions in the UMAP and number of groups in the HAC were chosen as the values that optimized the increase of acoustic group number while minimizing the decrease in mean and median silhouette values.

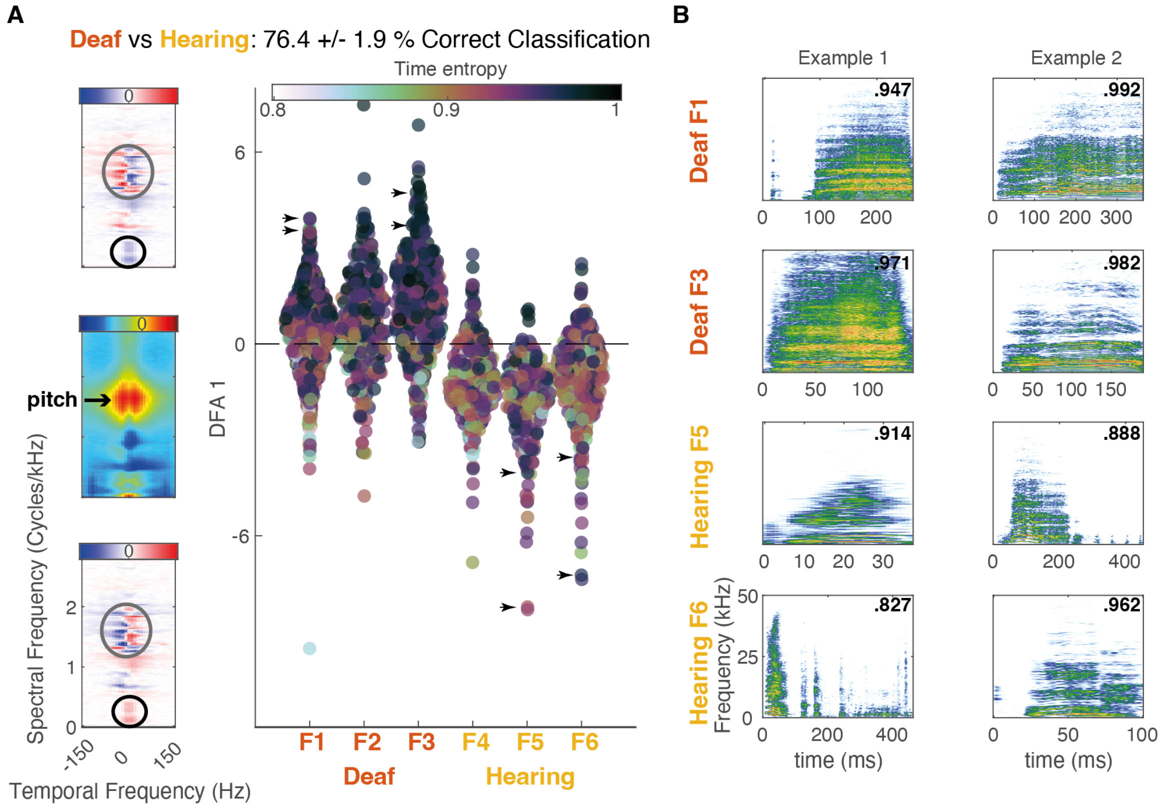


**Fig. S11.** Effect of deafness on female calls from Acoustic group 1.
A. For each female bat, projections of the vocalizations from acoustic group 1 on the linear acoustic axis that best discriminates between deaf (F1, F2 and F3) and hearing (F4, F5 and F6) bats. Each disk in the scatter plot represents a vocalization. The disk color indicates the value of entropy calculated on the amplitude envelope (Time Entropy) of the vocalization (high values indicate smoother calls). The three images on the left show the average MPS of vocalizations in Acoustic group 1 (middle) and the spectro-temporal changes in the MPS (discriminant function) that best characterize vocalizations from deaf female bats (top) and vocalizations from hearing female bats (bottom). Note that while on average female vocalizations in acoustic group 1 are characterized by pitch values of ~1.6c/kHz corresponding to ~700Hz (black arrow), the time varying pitch modulations are different between deaf and hearing females with a complex mixture of upsweeps and downsweeps depending on the value of the fundamental (gray circles). Deaf female calls were also characterized by a decrease in energy in the region of formants (black circles). The black arrows in the scatter plot point to the example calls shown in B.
B. Spectrogram of 2 example calls from acoustic group 1 by four female bats. Note that vocalizations from deaf bats are less variable in amplitude over time than vocalizations from hearing bats, leading to higher time entropy for deaf bats vocalizations. Values of time entropy are given for each example in the upper right corner of the spectrogram. The audio file of each of these examples can be found as supplementary sounds (from left to right and top to bottom: Sound S17-S24).

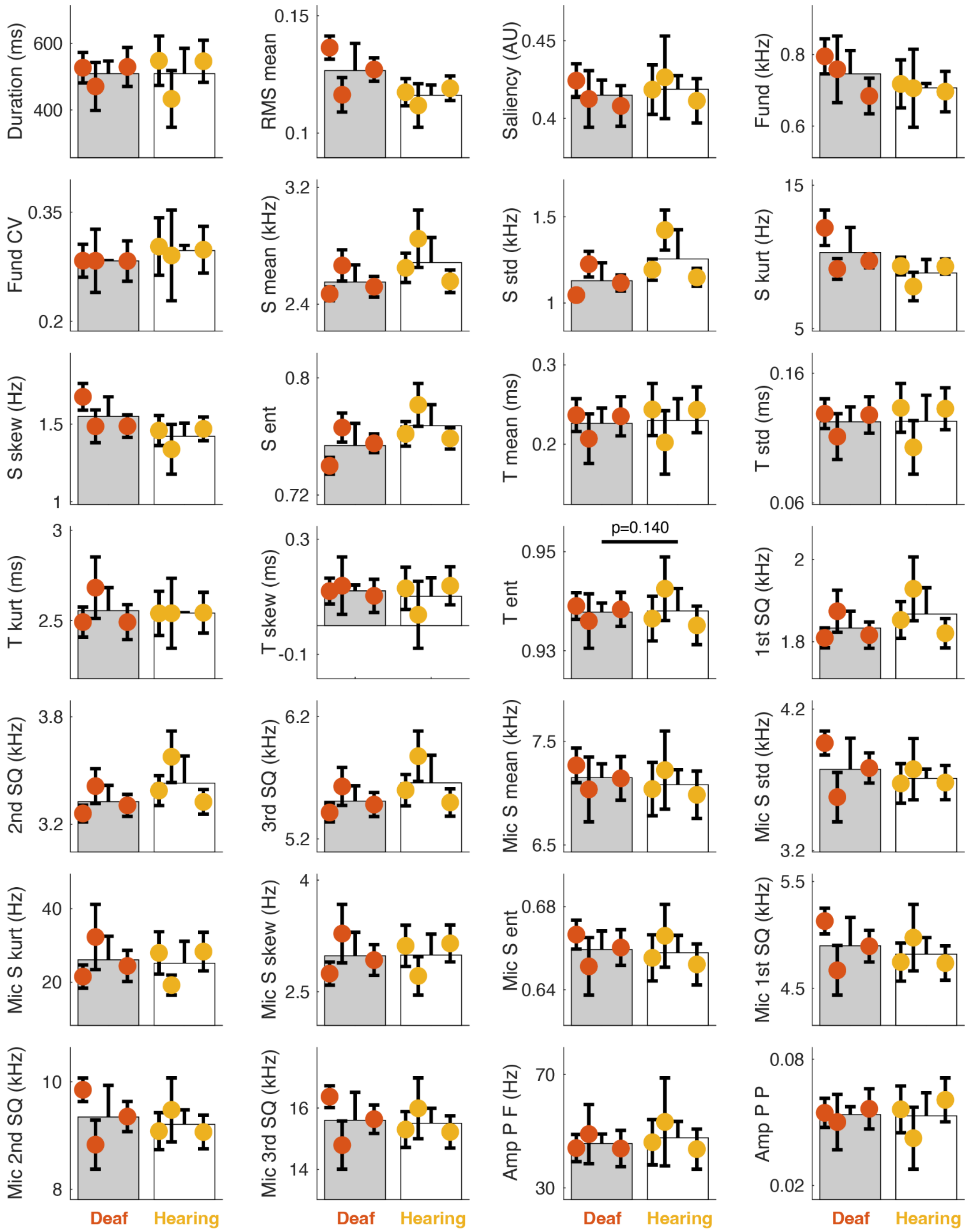


**Fig. S12.** Average acoustic features of female bats calls in Acoustic Group 1.
28 acoustic features were chosen to describe both the temporal and the spectral content of vocalizations (BioSound library, see definitions of acoustic features in legend of Figure S3). Average values for each bat are depicted by the dots color coded according to treatment. The bars depict the average values across deaf (gray) and hearing (white) bats. Error bars are ± 2*STE. Time entropy, a measure of vocalization rhythmicity, was the only significant ($p<0.01$) acoustic parameter between deaf and hearing bats in a the log likelihood ratio test comparing two Linear Mixed effect model with and without hearing ability as a predictor and with Bat identity as a

random variable. However, this parameter did not remain significant after FDR correction ($p>0.05$). P-values are given after FDR correction for acoustic features that were significant before FDR correction although they were calculated for all features.

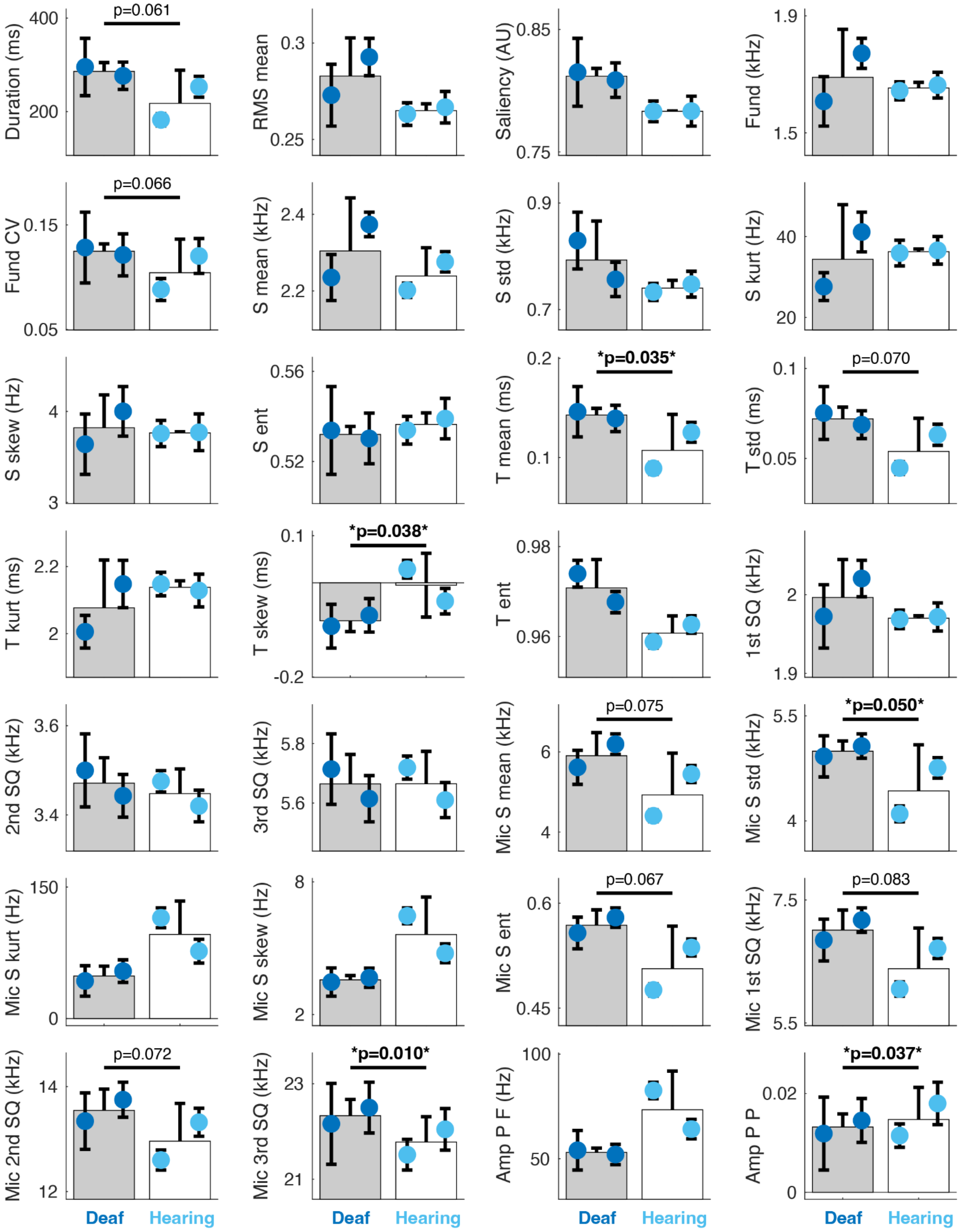


**Fig. S13.** Average acoustic features of male bat calls in Acoustic Group 4.
28 acoustic features were chosen to describe both the temporal and the spectral content of vocalizations (BioSound library, see definitions of acoustic features in legend of Figure S3). Average values for each bat are depicted by the dots color coded according to treatment. The bars depict the average values across deaf (gray) and hearing (white) bats. Error bars are ± 2*STE. 12/28 acoustic parameters were significantly different between deaf and hearing bats in a log likelihood ratio test comparing two Linear Mixed effect models with and without hearing ability

as a predictor and with Bat identity as a random variable ($p<0.01$, black lines). However, only five parameters remained significant after FDR correction (Time mean, Time skewness, Microphone Spectral standard deviation, Microphone third spectral quartile, and Amplitude periodicity Power). P-values are given after FDR correction for acoustic features that were significant before FDR correction although they were calculated for all features.

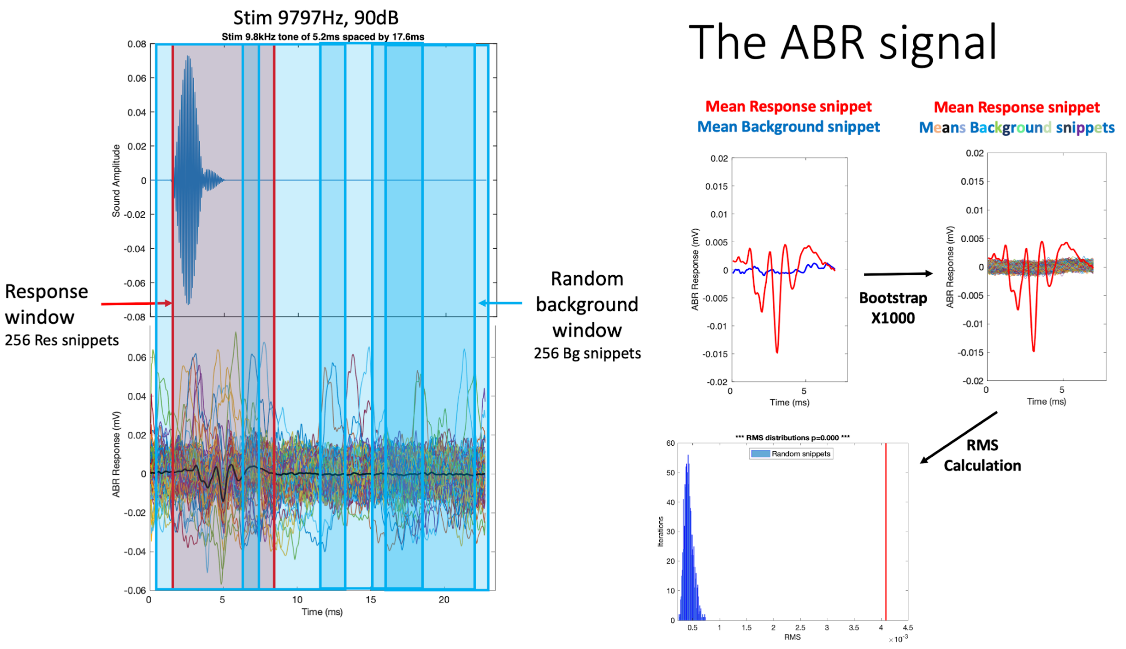


**Fig. S14.** ABR signal extraction and processing. See description in Audiogram section. Note that only four background windows are shown as examples in the figure, when there should be 256, one for each ABR signal recorded for each presentation of each stimulus. Bg, Background; Res, Response.

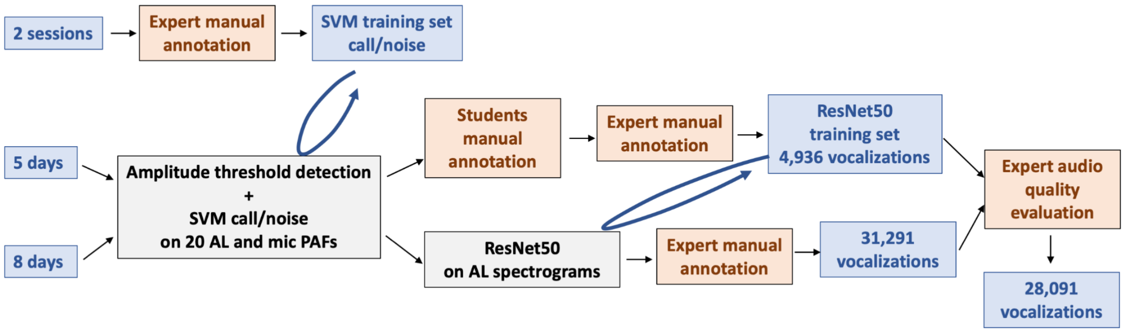


**Fig. S15.** Flow chart describing the vocalization discovery procedure.
AL, audiologger; mic, microphone; PAFs, predefined acoustic features.

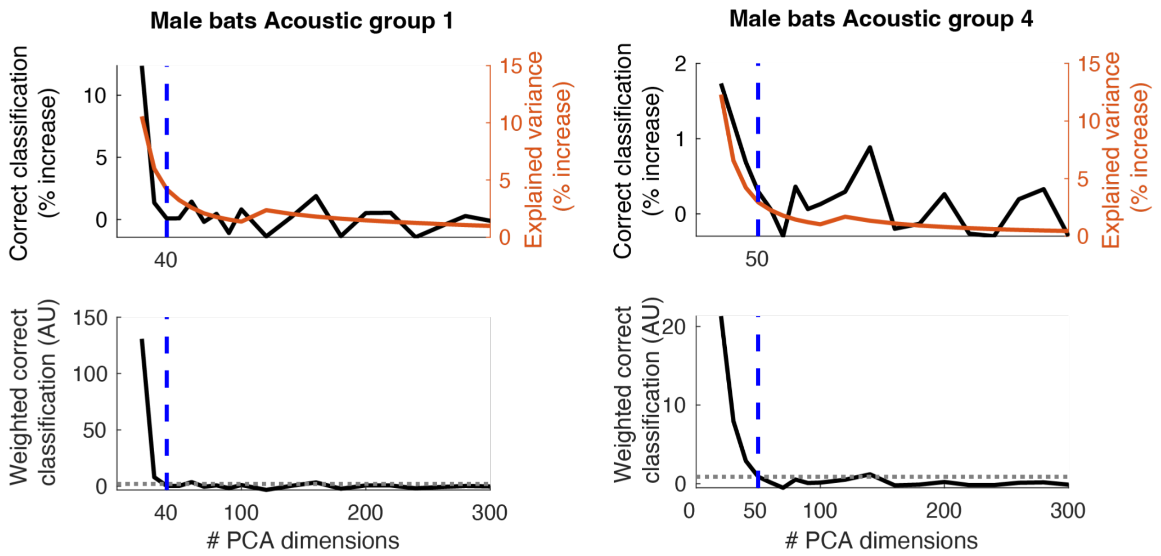


**Fig. S16.** Optimization of the number of dimensions in the Regularized permutation DFA for male calls.

For each acoustic group, the top plot displays the change in performance of classification of deaf vs hearing as the number of PCA dimensions increase (black line) and the percentage of explained variance that are brought by the added PCA dimensions compared to the previous step (red line). The bottom plot displays the noise threshold (dotted gray line, half of the average standard deviation of the classification performance over all tested PCA dimensions) and the measure used as a decision criteria: the ratio between the correct classification and the explained variance (ratio between the black and the red line in the top plot). In all plots, the optimal number of dimensions when the weighted correct classification drops below the noise level is indicated by the blue dotted line.

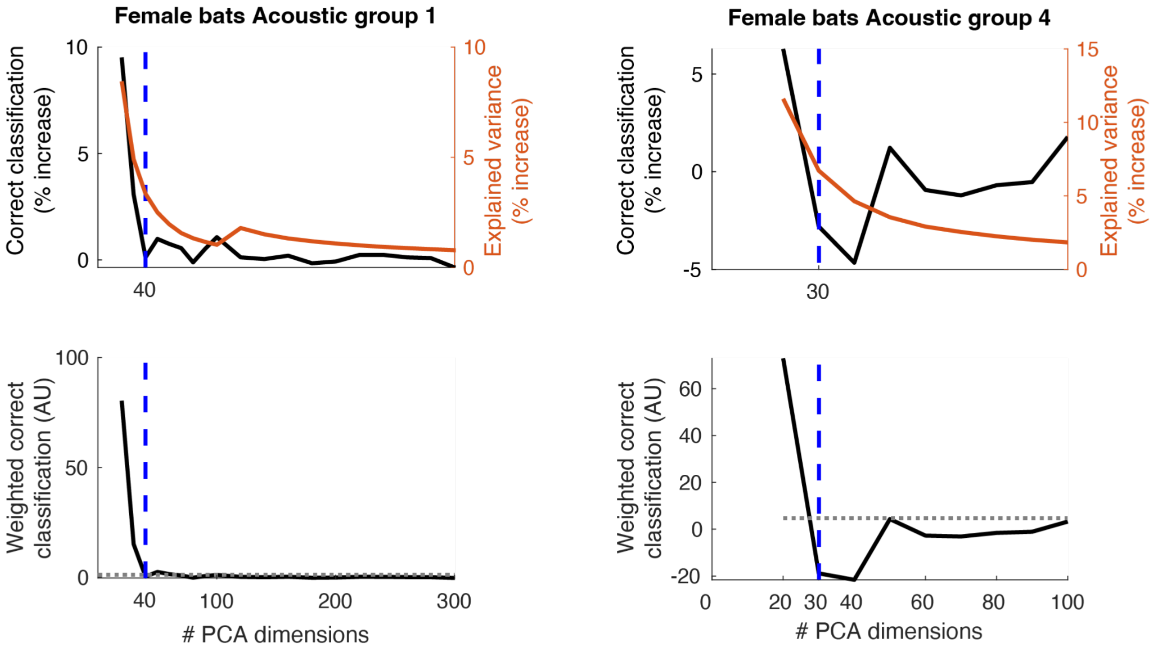


**Fig. S17.** Optimization of the number of dimensions in the Regularized permutation DFA for female calls.

For each acoustic group, the top plot displays the change in performance of classification of deaf vs hearing as the number of PCA dimensions increase (black line) and the percentage of explained variance that are brought by the added PCA dimensions compared to the previous step (red line). The bottom plot displays the noise threshold (dotted gray line, half of the average standard deviation of the classification performance over all tested PCA dimensions) and the measure used as a decision criterion: the ratio between the correct classification and the explained variance (ratio between the black and the red line in the top plot). In all plots, the optimal number of dimensions when the weighted correct classification drops below the noise level is indicated by the blue dotted line.

**Sound S1 (separate file).** Example Group 4 call from deaf male M2 recorded from ambient microphone and amplified to use the full dynamic range.

**Sound S2 (separate file).** Example Group 4 call from deaf male M2 recorded from ambient microphone and amplified to use the full dynamic range.

**Sound S3 (separate file).** Example Group 4 call from deaf male M1 recorded from ambient microphone and amplified to use the full dynamic range.

**Sound S4 (separate file).** Example Group 4 call from deaf male M1 recorded from ambient microphone and amplified to use the full dynamic range.

**Sound S5 (separate file).** Example Group 4 call from hearing male M3 recorded from ambient microphone and amplified to use the full dynamic range.

**Sound S6 (separate file).** Example Group 4 call from hearing male M3 recorded from ambient microphone and amplified to use the full dynamic range.

**Sound S7 (separate file).** Example Group 4 call from hearing male M4 recorded from ambient microphone and amplified to use the full dynamic range.

**Sound S8 (separate file).** Example Group 4 call from hearing male M4 recorded from ambient microphone and amplified to use the full dynamic range.

**Sound S9 (separate file).** Example Group 1 call from hearing female F4 recorded from ambient microphone and amplified to use the full dynamic range.

**Sound S10 (separate file).** Example Group 1 call from hearing male M3 recorded from ambient microphone and amplified to use the full dynamic range.

**Sound S11 (separate file).** Example Group 2 call from hearing male M3 recorded from ambient microphone and amplified to use the full dynamic range.

**Sound S12 (separate file).** Example Group 2 call from hearing male M3 recorded from ambient microphone and amplified to use the full dynamic range.

**Sound S13 (separate file).** Example Group 3 call from hearing female F4 recorded from ambient microphone and amplified to use the full dynamic range.

**Sound S14 (separate file).** Example Group 3 call from hearing male M4 recorded from ambient microphone and amplified to use the full dynamic range.

**Sound S15 (separate file).** Example Group 4 call from hearing male M4 recorded from ambient microphone and amplified to use the full dynamic range.

**Sound S16 (separate file).** Example Group 4 call from hearing male M3 recorded from ambient microphone and amplified to use the full dynamic range.

**Sound S17 (separate file).** Example Group 1 call from deaf female F1 recorded from ambient microphone and amplified to use the full dynamic range.

**Sound S18 (separate file).** Example Group 1 call from deaf female F1 recorded from ambient microphone and amplified to use the full dynamic range.

**Sound S19 (separate file).** Example Group 1 call from deaf female F3 recorded from ambient microphone and amplified to use the full dynamic range.

**Sound S20 (separate file).** Example Group 1 call from deaf female F3 recorded from ambient microphone and amplified to use the full dynamic range.

**Sound S21 (separate file).** Example Group 1 call from hearing female F5 recorded from ambient microphone and amplified to use the full dynamic range.

**Sound S22 (separate file).** Example Group 1 call from hearing female F5 recorded from ambient microphone and amplified to use the full dynamic range.

**Sound S23 (separate file).** Example Group 1 call from hearing female F6 recorded from ambient microphone and amplified to use the full dynamic range.

**Sound S24 (separate file).** Example Group 1 call from hearing female F6 recorded from ambient microphone and amplified to use the full dynamic range.
